# Fluid-Niche and Microglial Signatures Prime Robust Intraventricular Macrophage Response to Blood During Brain Development

**DOI:** 10.1101/2025.10.02.679906

**Authors:** Miriam E. Zawadzki, Christine Hehnly, Jordan C. Benson, Dario X. Figueroa Velez, Sivan Gelb, Ceva Stanley, Lillian I.J. Byer, Huixin Xu, Cameron Sadegh, Aja Pragana, Maria K. Lehtinen

## Abstract

Intraventricular macrophages (IVMs) reside in cerebrospinal fluid (CSF) and are considered a border-associated macrophage (BAM) population in the brain. Although they represent the first line of defense against intraventricular challenges, their developmental roles and responses to injury are poorly understood. This knowledge is relevant for conditions including neonatal intraventricular hemorrhage (IVH), where blood extravasates into brain ventricles, leading to life-long negative sequelae including cerebral palsy and hydrocephalus. Here, we show that IVMs are first responders to blood in developing brain ventricles, phagocytosing red blood cells and upregulating iron-processing machinery. Live imaging of developing mouse ventricles and choroid plexus revealed that IVMs are dynamic and morphologically distinct from non-IVM macrophages. Their transcriptional profiles distinguish them from other BAMs as they also exhibit signatures of “youth-associated microglia” and characteristics of cavity macrophages found in fluid niches such as the peritoneum. Our findings provide insights into IVM development and function, highlighting their therapeutic potential.

## INTRODUCTION

Maintaining a healthy cerebrospinal fluid (CSF) environment is critical for brain development and lifelong homeostasis. The choroid plexus (ChP) secretes and regulates the bulk of CSF contents in the brain ventricles and is believed to play a critical role in immune access^1–4^. Intraventricular macrophages (IVMs) found in the CSF and on the ChP are believed to scavenge the entire ventricular environment and to phagocytose and remove debris, pathogens, and unwanted damaged/dead cells.^5,6^ However, it is not clear whether IVMs significantly affect debris clearance, which would increase their potential utility as therapeutic targets that may influence clinical outcomes. Elucidating IVM function is particularly salient following neonatal intraventricular hemorrhage (IVH), a condition in which blood from ruptured vessels in the developing brain’s the germinal matrix flows into the brain’s lateral ventricles. Occurring in 25-30% of infants born earlier than 29 weeks gestation, long-term sequelae of IVH include cerebral palsy, developmental delay, and post-hemorrhagic hydrocephalus (PHH), defined by the dysregulation of CSF production and resorption.^7–10^ Ample data indicate significant neurotoxicity of iron and hemoglobin released from red blood cells (RBCs).^11–14^ Levels of CSF ferritin and hemoglobin in IVH patients positively correlate with the development and severity of PHH; a longitudinal decrease in CSF ferritin improved cognitive and motor outcomes.^15,16^ Moreover, intracerebroventricular (ICV) injections of iron alone is sufficient to cause PHH in animal models.^17^ Taken together, these data suggest that RBC removal efficiency is a critical process for brain health and that IVMs may play a role in this process.

Macrophages are often categorized by their ontogeny and location, including in the CNS.^18^ First described by Kolmer over one century ago, IVMs are found throughout the brain’s ventricular system and gather at the ChP (e.g., as ChP epiplexus cells).^19,20^ These tissue resident macrophages enter the ventricles shortly following neurulation and collectively patrol three important brain compartments: the ChP (blood-CSF barrier), the CSF, and the ventricular lining (brain-CSF barrier).^21^ Prior studies have begun to characterize the molecular identities of these cells.^5,22,23^ Here, using mouse models, we developed new tools to investigate the identity, movements, and functions of IVMs during brain development in the context of baseline surveillance and responses to intraventricular RBCs. Utilizing the stromal macrophages in the ChP to contrast to IVMs, we investigated their temporal dynamics across developmental stages and following administration of blood products into the ventricles. Our analyses defined three molecularly and anatomically distinct macrophage populations. Further, embryonic IVMs were plentiful, exhibited surveillance behavior, and could respond to RBCs in the CSF by clearing the cells and sequestering iron. These findings highlight a role for macrophages in the developing brain to facilitate recovery from IVH. Boosting this process may be an effective strategy to ameliorate brain damage and perhaps prevent PHH.

## RESULTS

### IVM abundance, morphology, and mobility are developmentally regulated

We identified that already early in brain development, epiplexus ChP macrophages are a distinct population from stromal ChP macrophages, with marked developmental shifts in abundance and morphology. Using brain sections, we quantified IVMs across three developmental timepoints: E14.5 (embryonic), P7 (neonatal), and adult (approximately 4 months) (**Figure 1A-C**) and found that epiplexus cells and total IVMs were 2-3 fold greater in embryonic brain ventricles relative to postnatal timepoints (**Figure 1D-E**). Since intact tissue provides clearer visualization of cell morphology than sectioned tissue, which disrupts structures during slicing, we used ChP explants to distinguish stromal and ventricular macrophages. Both stromal and epiplexus ChP macrophages are capable of phagocytosis of surrounding particles.^20^ We differentially labeled macrophages using spectrally-distinct large fluorescent dextrans delivered directly ICV or introduced into the peripheral circulation either through IP (neonates) or placental (embryos) injections in CX3CR1^GFP/+^ reporter mice (**Figure 1F-G**).^24^

**Figure 1.**
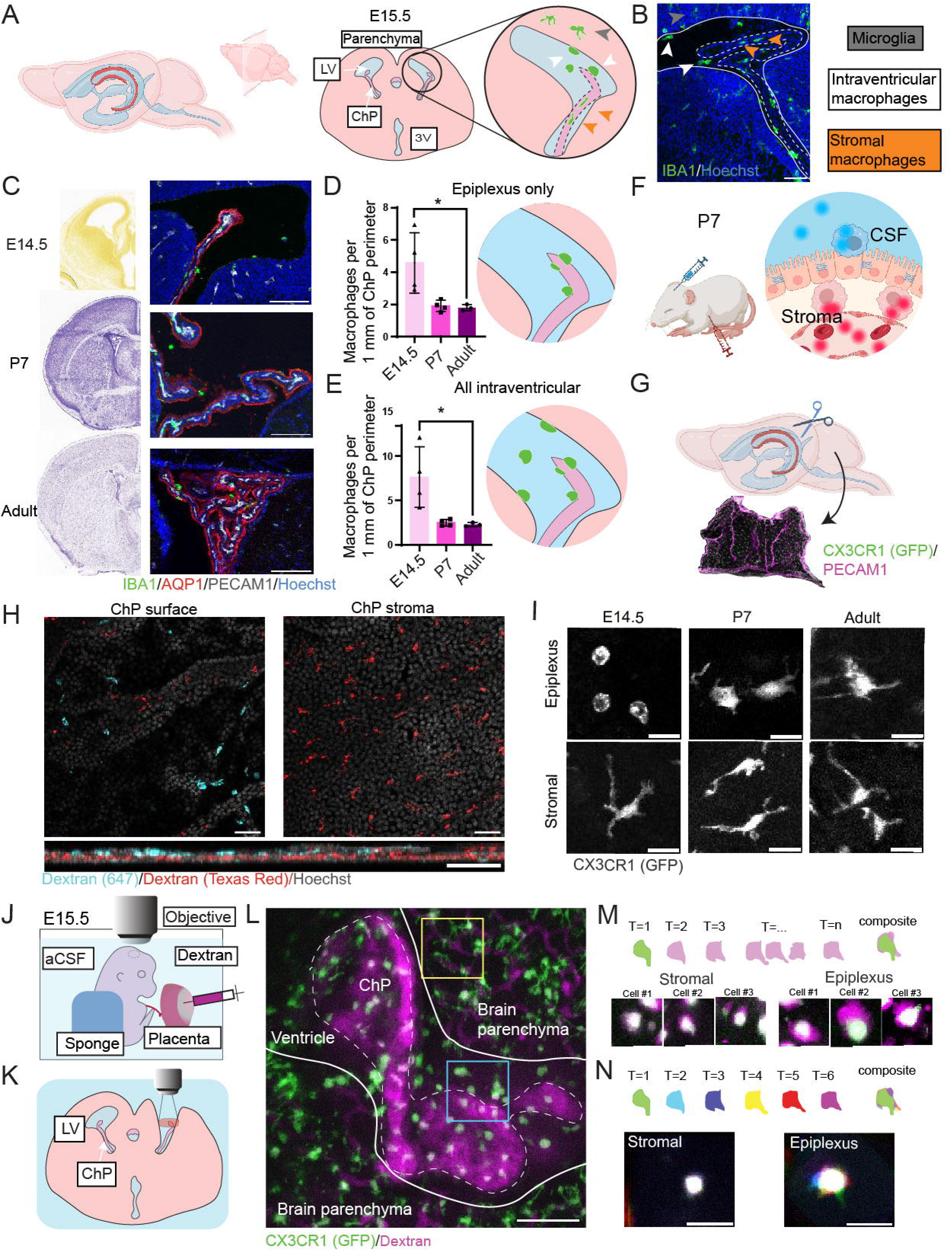
Epiplexus cells are distinct from their stromal counterparts morphologically and behaviorally throughout development. (**A**) Sagittal schematic of ventricles (blue) and lateral ChP (pink) in the developing mouse brain. LV= lateral ventricle, ChP = choroid plexus, 3V = third ventricle. Made using Biorender.com. (**B**) Coronal schematic and image of an E15.5 mouse ventricle showing microglia (gray), IVMs (white) and stromal macrophages (orange). Scale bar = 50 µm. (**C**) Coronal hemisections of mouse brains at E14.5, P7, and adult with matched ChP images. Allen Mouse Brain Atlas, <u>mouse.brain-map.org</u> and <u>atlas.brain-map.org</u>. Scale bar = 100 µm. (**D-E**) Quantification of density of epiplexus (D) and total IVM (E) density across ages. (Kruskal-Wallis with Dunn **p<0.01 for both, *p<0.05 for epiplexus and total IVM post-hoc multiple comparisons E14.5 vs adult). (**F-G**) Schematic of dual-colored dextran injections ICV and IP (F) and lateral ventricle ChP whole mount (G). Made using Biorender.com. (**H**) P7 ChP wholemount 3 hours following dextran injections, showing surface (left) and stromal (right) layers. Orthogonal view below. Scale bar = 50 µm. (**I**) Stromal and epiplexus macrophages at different ages CX3CR1^GFP/+^ animals, gray = GFP. Scale bar = 5 µm. (**J-K**) *Ex vivo* E15.5 embryo setup (J, Dotted circle indicates dissected cerebral cortex, aCSF= artificial CSF) and live imaging schematic (K). (**L**) Live 25x image of E15.5 embryonic ChP (dashed) and brain parenchyma (solid) with microglia (yellow box) and ChP macrophages (blue box) (see **Supplemental Videos S1 and S2**). Scale bar = 100 µm. (**M**) Schematic and representative stromal and epiplexus cell movement over 10 min, 45 sec from (L), averaging all movement (purple) compared with starting frame (green), see **Supplemental Video S3.** (**N**) Additional schematic and representative stromal and epiplexus cell movement pseudocolored by sampled timepoint (6 total timepoints sampled throughout 10 minutes of video, see **Supplemental Video S3**).

Three hours later, histology revealed distinct labeling patterns, consistent with these macrophages surveilling distinct fluid compartments. Stromal macrophages (red dextran) displayed ramified processes across development, whereas epiplexus cells (far-red dextran, in blue) appeared largely devoid of processes and displayed a rounded morphology that became more ramified in adulthood (**Figure 1H-I, Supplemental Figure S1A-B**).

To characterize macrophage behavior in living tissue, we adapted our two-photon imaging platform to study E15.5 mouse ChP *ex vivo* (**Figure 1J-K**).^24–26^ We used the lateral ventricle ChP blood vessel topography to establish the anatomical location of macrophages with respect to neighboring brain parenchyma and ventricle walls (**Figure 1L**, **Supplemental Video S1**). Microglia in the brain parenchyma were branched and ramified with long processes that continuously extended and retracted.^27,28^ By contrast, the ChP macrophages were relatively ameboid (**Supplemental Video S2**).^25^ ChP macrophage location could be identified by dextran uptake and perivascular localization (**Supplemental Figure S1C**). Two-photon imaging showed stromal cells were relatively immobile; however, short epiplexus processes were highly motile with active extension and retraction (**Figure 1M-N; Supplemental Video S3**).^25^ This behavior of ChP epiplexus macrophages suggests they engage in local sampling and surveillance. Taken together, these findings demonstrate that embryonic ChP epiplexus macrophages are distinct in their abundance, amoeboid morphology, and heightened motility.

### Effects of RBC injection on IVM morphology, abundance, and activity

To observe embryonic epiplexus macrophages in the context of blood exposure and iron sequestration in the CSF, we adapted our established model of intraventricular hemorrhage by using ICV RBC delivery (**Figure 2A-B**)^9^. After 24 hours, IVMs contained phagocytosed RBCs, displayed amoeboid and vacuolated morphologies, and were more abundant (**Figure 2C-F**). This expansion was not reduced in *Ccr2*-deficient (CCR2^RFP/+^) and *Ccr2*-knockout (CCR2^RFP/RFP^) mice (**Figure 2G**). In *Ccr2^RFP/+^* embryos, RBC injection rarely yielded RFP^+^ and IBA1^+^ macrophages at 24 hours, indicating minimal peripheral monocyte contribution to IVM expansion (**Figure 2H**). Furthermore, two-photon imaging (**Figure 2I-J**) revealed somatic translocation and rapid process dynamics in RBC-exposed macrophages compared to saline-injected controls, suggesting the model that resident cells themselves mount the RBC-induced response (**Supplemental Video S4**).

**Figure 2.**
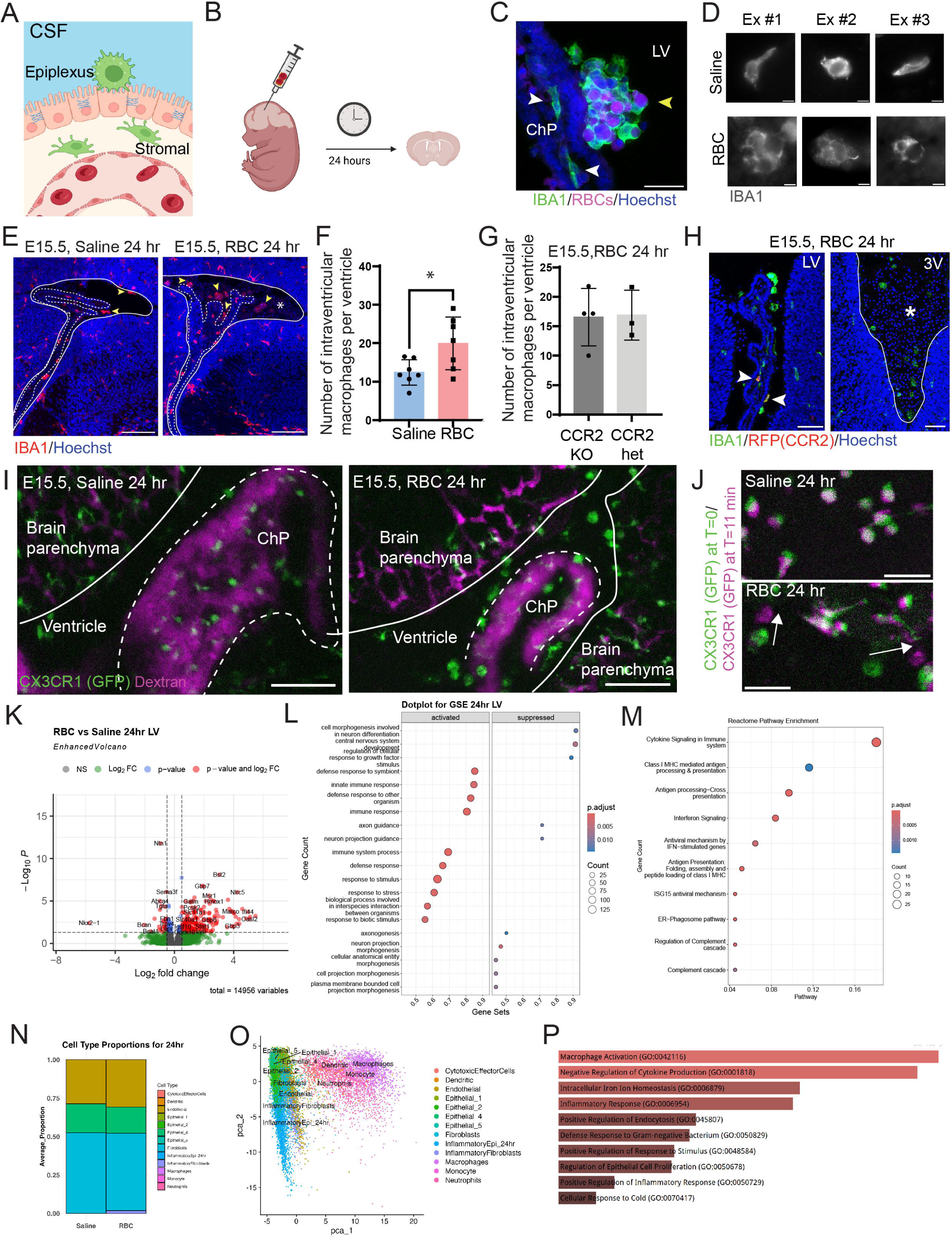
Robust IVM response to blood exposure. (**A**) Cross-sectional schematic of epiplexus and stromal cell locations. Made using Biorender.com. (**B**) Injection and collection timeline. (**C**) A cluster of epiplexus macrophages containing whole RBCs. Scale bar = 25 µm. **(D)** Macrophage (IBA1) staining 24 hours following saline or RBC injection. Scale bar = 5 µm. (**E-F**) Representative tiled 10x images 24 hours following injection showing localization and immunoreactive macrophages (E, IBA1, yellow arrowheads. Scale bar = 100 µm. Asterisk denotes RBCs nuclei.) with quantification of IVMs (F, *p < 0.05, Welch’s t-test). (**G-H**) quantification (G, Welch’s t-test p = 0.9242) and representative images (H, arrowheads = IBA1+, CCR2+. Scale bar = 50 µm. Asterisk denotes RBCs nuclei) 24 hours following ICV RBCs in *Ccr2*^RFP/+^ or *Ccr2*^RFP/RFP^ embryos. (**I**) *Ex vivo* E15.5 ChP (dashed) and parenchyma (solid) 24 hours following saline or RBC ICV delivery. Contrast adjusted for vessel visualization. Scale bar = 100 µm. (**J**) Representative images from **Supplemental Video S4**. Scale bar = 50 µm. (**K**) Volcano plot of ChP RNA Seq. (**L-M**) GO terms (L) and Reactome (M) pathway enrichment of these terms. (**N-O**) Predicted cell proportion (N) and identities (O) from deconvolution after RBC injection. (**P**) GO terms for upregulated macrophage genes.

Having established that RBCs increase IVM number and activity independent of monocyte recruitment, we next examined transcriptional programs underlying this response. We used bulk RNA sequencing of pooled whole ChP tissue from embryos 24 hours following saline or RBC injection and found that RBC exposure triggers an interferon-driven inflammatory response in the ChP and activates iron-clearance pathways in ChP macrophages (**Supplementary Data Table 1**). Differential expression identified upregulation of immune activation including *Nfkb1*, *Irf7*, *Socs3*, and *Cxcl10* (**Figure 2K, L**). Interferon-stimulated genes including *Irf7*, *Ifit1*, and *Isg15*, suggested activation of viral-sensing pathways following nucleated embryonic RBC exposure (**Figure 2M**).^29^ Downregulated genes instead mapped to neurodevelopmental pathways, including neuronal differentiation (*Neurod6*, *Tubb3*), axon guidance (*Sema3c*, *Dcc*), and synaptic organization (*Nlgn1*, *Grin1*), suggesting that RBC-induced inflammation consistent with the suppression of neuroglia-like cells previously described in E16.5 ChP.^30^ Deconvolution using single cell data from inflammatory ChP collected from LPS treated mice, revealed increased macrophage signatures in RBC samples (**Figure 2N**).^31^ UMAP-based deconvolution using differentially expressed genes, further resolved activated macrophages, dendritic cells, and monocytes with *Irf7*, *Socs3*, and *Cxcl10* enriched in these subsets (**Figure 2O**). In RBC-injected compared to saline cohorts, macrophage-associated signatures showed enhanced transcription of genes linked to activation, inflammatory response, and intracellular iron homeostasis (**Figure 2P**).

Consistent with their phagocytosing RBCs, upregulated macrophage genes included *Hmox1* and *Slc11a1*, key regulators of heme metabolism and iron transport respectively^32,33^. We confirmed *Hmox1* and *Slc11a1* expression increase using qPCR and found that HMOX1 protein predominated in IVMs using IHC (**Figure 3A-B, Supplemental Figure S1D-E**). Given the upregulation of iron-handling genes, we expected IVMs to maintain significant iron stores. Indeed, days following IVH in *Cx3cr1*^GFP/+^ animals, we observed iron sequestration solely in GFP-positive cells in and around the ventricle by Perl’s Prussian blue staining for hemosiderin (**Figure 3C-D**).

**Figure 3.**
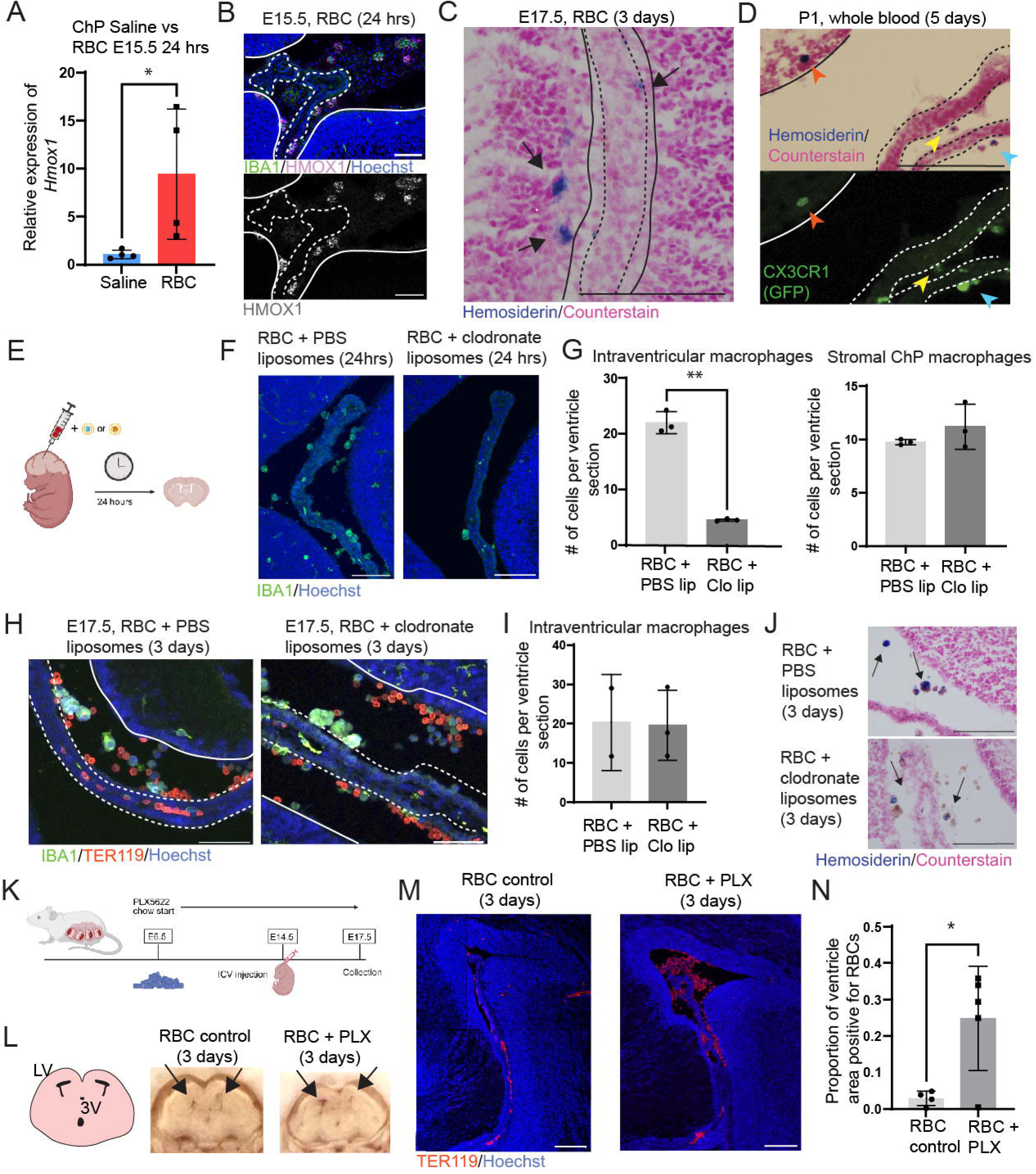
IVMs persist during hemorrhage and contribute to clearance of RBCs from the ventricle. (**A-B**) *Hmox1* qPCR (C, Mann-Whitney *p <0.05) and Representative HMOX1 staining in ChP 24 hours RBC injection. Scale bar = 50 µm (**C**) Representative of ChP (dashed) and ventricle (solid) Perls stain of hemosiderin cells (black arrow) 3 days post-RBC injection. Scale bar = 100 µm. (**D**) Representative images 5 days following whole blood injection of ChP (dashed) and parenchyma (solid) of hemosiderin cells (Perls, top) and macrophage (CX3CR1^GFP/+^, bottom) with overlapping cells color coded by arrowhead. Scale bar = 100 µm. **(E)** Schematic of experimental protocol for ICV delivery of RBCs with liposomes containing PBS (blue) or clodronate (orange). Made using Biorender.com. (**F-G**) Representative image of macrophages (IBA1) in lateral ventricle (F) and quantification of IVMs and stromal macrophages 24 hours following injection (G, **p<0.005 (p=0.0039) and p=0.3589, respectively, Welch’s t-test) (**H-I**) Representative image of macrophages (IBA1^+^) in lateral ventricle (H) and quantification (I) of IVMs 3 days following injection (p=0.9484, Welch’s t-test). Scale bar = 100 µm. (**K**) Experimental schematic for PLX5622 administration and macrophage depletion in embryos. Made using Biorender.com. (**L-N**) Gross brain image (L), RBC (TER119) staining, scale bar = 150 µm (M) and quantification of RBCs (N) with and without exposure to PLX5622 (*p<0.05, p= 0.0254, Welch’s t-test).

### IVMs are necessary to clear RBCs from the embryonic ventricle

To determine the necessity of IVMs in clearing RBCs, we first tested ICV-injected clodronate liposomes.^34^ ICV clodronate liposomes depleted IVMs specifically by 24 hours, but the cells repopulated within 3 days and were indistinguishable from controls in the number of macrophages and in iron sequestration function (**Figure 3E-J**). Therefore, we next depleted total myeloid cells, including IVMs, by treating pregnant dams with CSF1R inhibitor, PLX5622.^35,36^ Three days following RBC injection in myeloid-depleted mice, no IVMs remained in PLX5622-treated embryos, and their ventricles retained significantly more RBCs than controls with intact IVMs (**Figure 3K-N**). Thus, initial phagocytosis by IVMs may substantially contribute to RBC clearance from the ventricle three days after hemorrhage, consistent with their necessity for this process in the embryonic brain.

### IVMs and stromal ChP macrophages have distinct transcriptional identities

Embryonic ChP macrophages segregated into transcriptionally and functionally distinct populations aligned with their local environments. Analysis of our previously generated single-cell RNA sequencing (scRNA-Seq) atlas of E16.5 ChP revealed four clusters: *SpiC*+*Hmox1* (33.9%), *Lyve1*+*Cd163* (29.4%), *Cd9*+*Spp1*+*Gpnmb* (18.8%), and a proliferative *Nusap*+*Ki67* cluster (17.9%) (**Figure 4A, Supplemental Figure S2A**).^30^ Gene ontology analysis of non-replicating clusters suggested distinct functions linked to their dominant expression programs (**Supplemental Figure S2B**). Using CellChat, we inferred potential communication between macrophages and epithelial, mesenchymal, and endothelial populations.^37,38^ All three populations had greater outgoing than incoming interaction strength, suggesting that macrophages may be primarily recipients of signals from their local niche (**Figure 4B-C**).

**Figure 4.**
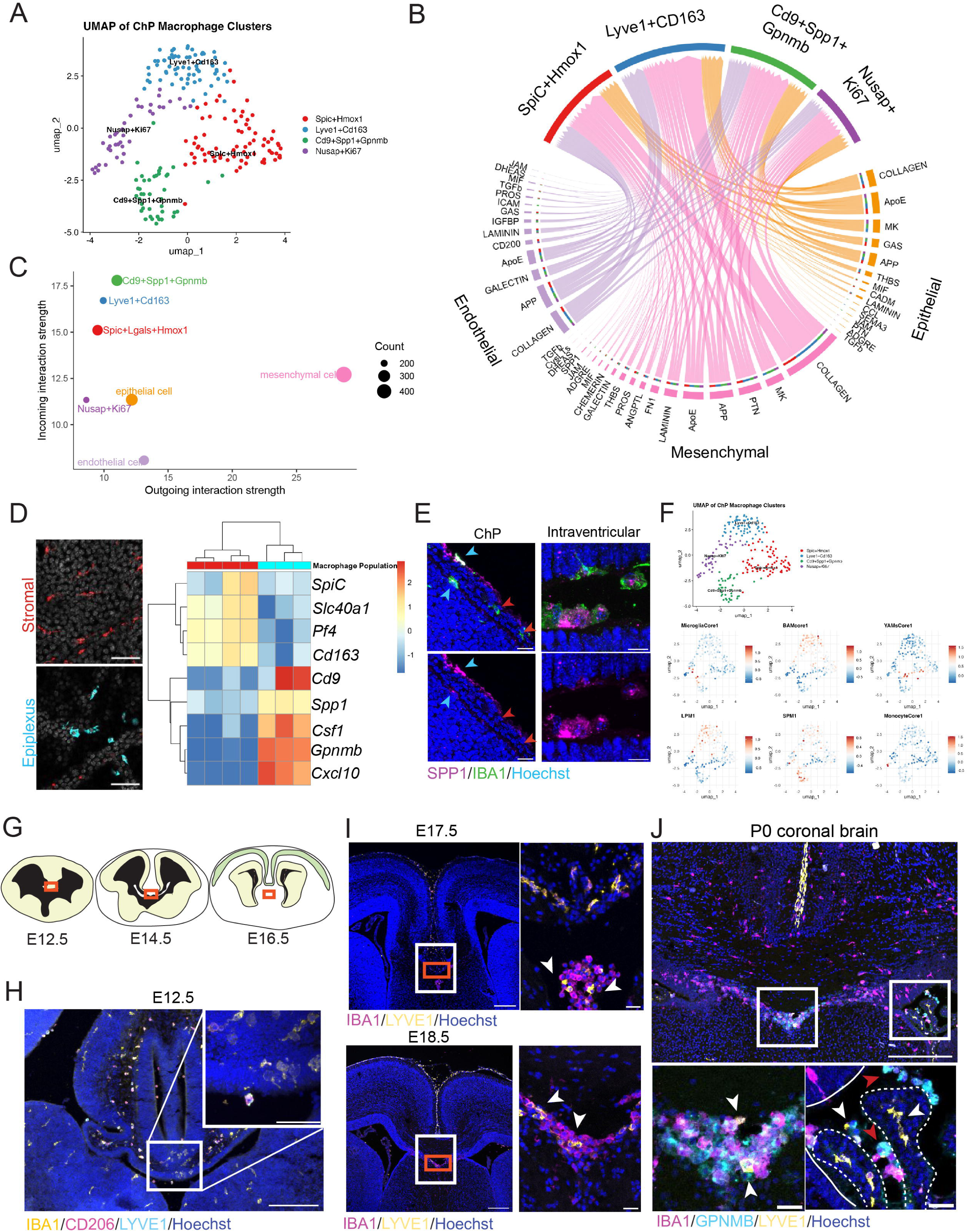
IVMs exhibit transcriptional signatures shaped by developmental origin and fluid niche environment, including ‘youth-associated’ microglial and peritoneal macrophage populations. (**A**) Myeloid cell population clustering from E16.5 ChP. (**B**) CellChat visualization of inferred outgoing signals from epithelial, mesenchymal, and endothelial cells to ChP macrophages. (**C**) Incoming vs. outgoing signal strength across populations. (**D**) Heatmap of selected differentially expressed genes from bulk RNA sequencing of sorted P7 stromal vs epiplexus macrophages. (**E**) Representative image of an E14.5 brain sections stained with SPP1. Scale bar = 10 µm. (**F**) UMAP of other myeloid population signatures reflected in the various ChP macrophage populations. (**G**) Schematic of choroid plaque development from E12.5-E16.5. (**H**) Representative image of an E12.5 brain section. Scale bar = 200 µm. Inset: Magnified view of choroid plaque. Scale bar = 50 µm. (**I**) Representative image of an E17.5 and E18.5 brain section showing the residual choroid plaque and underlying macrophages. Scale bar = 250 µm; 25 µm for inset. (**J**) Representative image of a P0 brain section showing the roughly equivalent region following the crossing of the corpus callosum as well as the lateral ventricle and ChP. Scale bar = 250 µm; 25 µm for inset.

We sought to understand the relationship between macrophage transcriptional signatures and anatomical location. First, we used ImageStream to confirm that following ICV dextran injection, epiplexus cells could be isolated (**Supplemental figure S1F-G**). Next, we performed dual dextran injections, isolated epiplexus and stromal macrophages by flow cytometry, and performed bulk RNA sequencing (**Figure 4D**, **Supplemental Figure S2C, Supplemental Data Table 2**). Hierarchical clustering showed that epiplexus cells upregulated genes consistent with the *Cd9*+*Spp1*+*Gpnmb* cluster, whereas stromal macrophages aligned with *SpiC*+*Hmox1* and *Lyve1*+*Cd163* clusters from our E16.5 dataset (**Figure 4D**). Immunostaining for SPP1, GPNMB, and CSF1 in E14.5 LV confirmed expression in intraventricular macrophages, including epiplexus cells, but not stromal macrophages (**Figure 4E, Supplemental Figure S2D**). Together, these findings suggest that IVMs, regardless of ventricular location, share a common transcriptional profile, consistent with a shared ontogeny and origin.

### IVMs share properties with YAMS and peritoneal macrophages

The transcriptional heterogeneity of ChP macrophages is shaped by both tissue origin and environment. To contextualize their program in a fluid-facing niche while also being resident brain macrophages, we compared epiplexus and stromal macrophages to canonical brain and peritoneal macrophages. At E16.5, our single cell RNA-seq revealed that epiplexus macrophages were enriched for youth-associated microglia (YAM) and canonical microglial signatures, while partially overlapping with small peritoneal macrophages (SPM) through expression of pro-inflammatory and sentinel genes (**Figure 4F, Supplemental Figure S2E-F**, Kruskal–Wallis p < 0.001). SPMs are monocyte-derived and epiplexus cells are known to be yolk-sac derived; their overlapping signatures did not include canonical monocytic genes^39,22,40,41^. In contrast, stromal macrophages aligned with BAMs and large peritoneal macrophages (LPM), expressing high levels of *Mrc1* and *Lyve1* while exhibiting comparatively low expression of classical LPM genes such as *Gata6* and *Timd4*, consistent with their embedding in the vascularized stroma but maintaining interactions with fluid environment of the blood. Bulk RNA-seq at P7 confirmed this divergence, with stromal macrophages retaining BAM-like programs and epiplexus cells preserving YAM-associated transcriptional features (**Supplemental Figure S2G**). YAMs, also known as axon-tract microglia (ATMs), are microglia that emerge beneath the corpus callosum and later migrate laterally across the ventricles, seen only in development.^42–44^ They canonically express *Gpnmb*, *Spp1*, *Csf1*, *Itgax*, and *Igf1.*^44^ In contrast to the YAMs, IVMs additionally expressed border-associated macrophage markers, such as CD206 (**Figure 4H**). We looked earlier in development to see if we could identify IVM origin at E12.5 and found that some macrophages present in the roof plate co-expressed markers including LYVE1 (a meningeal macrophage marker) and GPNMB (**Supplemental Figure S2H-I**). Recent evidence suggests that early in development some IVMs come from the choroid plaque, a structure from which the ChP develops at the ventricle edge of the roof plate.^21,45^ Indeed, we observed a population of roof plate macrophages appearing to breech the choroid plaque at E12.5 (**Figure 4H**).

Because of marker overlap, we considered the possibility that both IVMs and YAMs originate from this LYVE1+, GPNMB+ population in the roof plate. The macrophage-dense choroid plaque connects with the septum below to separate the two lateral ventricles between E12.5 and E15.5 (**Figure 4G**). The axon tracts of the corpus callosum begin to cross in late gestation above this region. This area at the midline is the cortico-septal boundary (CSB), also known as the cavum septum pellucidum (CSP), where ameboid microglia first appear in late embryonic ages in rodents.^46,47^ In concordance with this hypothesis, several of the macrophages in the CSP at E16.5/17.5 remained positive for LYVE1 despite having no clear residual anatomical connection to the meninges (**Figure 4I**). We also observed LYVE1 expression in some, but not all IVMs by birth (**Figure 4J**). The anatomical location of these macrophages/microglia, as well as the overlap of traditional ‘BAM’ and ‘microglial’ markers in the E12.5 yolk sac-derived myeloid cells, provide additional insight into how these two populations may be intimately related through their shared origin at the roof plate and choroid plaque.

## DISCUSSION

Here we show that epiplexus macrophages in the embryonic brain are plentiful, significantly more abundant than adult populations, and play an essential role in responses to foreign substances in the CSF as evidenced by phagocytosis of RBCs. Our newly-adapted imaging approaches paired with functional labeling and molecular analytic techniques provided a panoramic view of embryonic epiplexus cell responses to, and effects on, RBC presence in the immature ventricle, a condition that often leads to life-threatening and life-long brain damage.

Historically, the field has described epiplexus cells by histology and ultrastructural studies.^20^ Using real-time imaging, we directly visualized dynamic features of these cells including process motility that are otherwise impossible to obtain. This approach transforms a once-descriptive category into a functional framework. Epiplexus cell morphology and motility behavior was suggestive of local sampling and surveillance behavior, consistent with our previous live imaging observations of 4^th^ ventricle ChP (**Figure 1M-N, Supplemental Video S2**).^25^ Stromal counterparts, by contrast, displayed more ramified morphology and engaged directly with the nascent vasculature sampling the blood environment, reminiscent of our previous work in the lateral ganglionic eminence (synonymous to human germinal matrix), where vascular fragility underlies susceptibility to hemorrhage in prematurity.^26^ Together, these observations link form, function, and environment, revealing that the morphological heterogeneity of ChP macrophages aligns with distinct ecological niches.

Patterns of gene expression further enabled us to move from a descriptive imaging in fixed tissue to functional and developmental insights. Transcriptomic analysis from adult humans CSF myeloid cells span monocytic, BAM and microglial states, while newborn CSF is enriched in myeloid cells.^6,48–50^ Early fetal IVMs emerge during brain seeding by the yolk sac microglia.^51^ In adult mice, epiplexus macrophages cluster separately from other populations and display ‘microglial-like,’ signatures, however this has never been tested orthogonally.^5,6,22^ Our transcriptional analysis shows high expression of *Cd9*, *Gpnmb,* and *Spp1* in epiplexus cells consistent with phagocytic and tissue remodeling role of epiplexus cells, demonstrating a strong overlap with previously-demonstrated YAM programs.^31,47^ Immature microglia in developing human brains express activated microglial markers such as *Spp1* and *Apoe*, mirroring mouse YAM populations.^52^ YAMs/IVMs share a number of unique markers with adult damage-associated microglia (DAMs), suggesting that early brain environments requires these active macrophages.^53,42,54,55^ We further found that IVMs appear to originate from the midline choroid plaque, where YAMs appear days later, suggesting that they are the same developmental population in different environments.^45,47^ Infiltration from the midline is not limited to early development; recent evidence has shown that immune cells can track into the ventricles postnatally through the velum interpositum, a more posterior midline meningeal structure above the third ventricle.^56^ IVMs are present prior to the emergence of the ChP and their epithelial position suggests they may arrive via CSF, parallelling other macrophages trafficking through fluid niches.^31,57^ We found that epiplexus cells also overlap with SPMs, though unlike monocyte-derived SPMs, IVMs are yolk-sac-derived, highlighting that conserved sentinel programs can rise from distinct lineages.^22,40^ Together these findings define IVMs as yolk sac-derived sentinels at the CSF-brain interface, bridging developmental origin, morphological dynamics, and transcriptional identity with conserved role in phagocytic surveillance akin to YAMs.

As surveyors, IVMs are primed to respond to hemorrhage by phagocytosing RBCs, upregulating iron-processing genes and storing iron. Macrophage depletion from the whole brain, including IVMs, increased ventricular RBC burden and underscored their protective roles. In humans and preclinical models, clearance of blood products ameliorates post-hemorrhagic damage and hydrocephalus, a process likely critical in neonates where CSF outflow routes are immature.^58–62^ The meningeal lymphatics that serve as the main CSF outflow pathway develop in rodent models over the first month of life.^63,64^ Macrophages also mediate clot resolution after intraparenchymal hemorrhage, serving as a ‘egress route’ for RBCs and iron.^65,66^ The ChP also may serve as an iron removal route, as it expresses apical iron transporters, but intact RBCs cannot cross its tight junctions, requiring lysis or macrophage breakdown.^67,68^ Notably, protective phagocytosis must be balanced against inflammatory responses to hemorrhage, with studies showing conflicting effects on hydrocephalus following hemorrhage in the presence or absence of macrophages.^13,34,69,70^ Future studies are needed to define CSF clearance mechanisms and the baseline myeloid landscape in human neonates to enable therapeutic targeting of RBC update while limiting secondary inflammatory damage.

### Limitations of the study

Further dissecting IVM function will require new genetic tools that specifically ablate or block key gene programs. In our study, selective removal was not possible beyond 24 hours after RBC injection. PLX5622 eliminated IVMs, but also decimated other CNS macrophage populations, limiting attribution of RBC burden to IVM loss specifically. *Ex vivo* live imaging confirmed IVM motility in the lateral ventricle though invasive approaches may also have triggered cortical damage responses. Noninvasive modalities, such as three-photon imaging, could enable deeper, more physiological visualization.

## ACKNOWLEDGEMENTS

We thank members of the Lehtinen lab, Isaac Chiu, Susan Dymecki, Mark Fleming, and David Feliciano for helpful discussions and Nancy Chamberlin for critical reading and editing of the manuscript. We thank the following facilities and personnel: Chinfei Chen, Hisashi Umemori, Cheng-Hao Chien and the BCH IDDRC Cellular Imaging Core; Michael Anderson, Bin Bao, and the Cell Function and Imaging Core (Harvard Digestive Diseases Center); BCH: PCMM flow cytometry facility; the Broad Institute; Novogene for tissue processing and bulk-RNA sequencing. We thank Milo Taylor, Andrea Yessaillian, and Bradford Grant for technical expertise. We acknowledge https://BioRender.com for assistance with creating several schematics. This work was supported by: National Institutes of General Medical Sciences (NIGMS/NIH) grants T32 GM007753, T32 GM144273, and American Heart Association Pre-doctoral Fellowship (M.E.Z.); The Little Giraffe Foundation Neonatal Research Award 2021 (M.E.Z. and M.K.L.); NIH T32 NS007473, F32 NS136267, and Reagan Sloane Shanley Scholarship (C.H.); NIH NS129823-02S1 (J.C.B.); T32 NS007473, 1F32MH132178 (D.X.F.V.); Edward R. and Anne G. Lefler Center Postdoctoral Fellowship and Hebrew University Postdoctoral Fellowship (S.G.); BrightFocus postdoctoral fellowship in Alzheimer’s Disease research, BCH OFD Career Development Award (H.X.); Hydrocephalus Association, Pediatric Hydrocephalus Foundation, NIH R01 NS129823, RF1048790, The American Heart Association Innovator Award 23IPA1051611 (M.K.L.); BCH IDDRC 1U54HD090255. The content is solely the responsibility of the authors and does not necessarily represent the official views of the National Institute of General Medical Sciences or the National Institutes of Health.

## AUTHOR CONTRIBUTIONS

M.E.Z. and M.K.L. conceptualized and designed the study; M.E.Z., C.H., D.X.F.V. and M.K.L. established methodology; M.E.Z., C.H., J.C.B., S.G., H.X., and C. Sadegh. conducted experiments; M.E.Z., C.H., S.G., J.C.B., C. Stanley, A.P., and L.I.J.B. analyzed data; M.K.L. provided funding; M.K.L. supervised the study; M.E.Z., C.H., and M.K.L. wrote the manuscript. All co-authors read and approved the manuscript.

## RESOURCE AVAILABILITY

### Lead contact

Further information and requests for resources and reagents should be directed to and will be fulfilled by the Lead Contact, Maria Lehtinen

### Materials availability

This study did not generate mouse lines, viruses/plasmids, or other biological tools.

### Data and code availability

Code used for the registration of 2P videos can be found at https://github.com/LehtinenLab/Shipley2020 and HyperStackReg https://doi.org/10.5281/zenodo.2252521). Original data are available from the corresponding author upon request. Sequences are available on SRA BioProject PRJNA1332909 (ChP) and PRJNA1333390 (macrophages). All code is available at https://github.com/christinehehnly/MZ_CH_IVMs.git

## EXPERIMENTAL MODEL AND SUBJECT DETAILS

### Animals

All mouse work was performed in accordance with the Institutional Animal Care and Use Committees (IACUC) regulations and oversight at Boston Children’s Hospital. Eight-to ten-week-old CD1 breeders were purchased from Charles River Laboratories (CRL) and experiments used in-house bred timed-pregnant CD1 for E14.5 or P7 offspring. Embryonic live-imaging experiments: *Cx3cr1^gfp/gfp^* (RRID:IMSR_JAX:005582) were crossed with CD1 (RRID:IMSR_CRL:022) wild type female mice for the embryonic live imaging. Offspring were also used for E14.5/P7/adult explants following ICV and IP/intraplacental (E14.5 only) dextran injections. *Ccr2* KO experiment: *Ccr2^rfp/rfp^* (RRID:IMSR_JAX:017586) males were crossed with *Ccr2 ^rfp/+^* females to generate offspring. RFP/RFP animals were compared to their RFP/+ littermates. All animals were housed under 12hr day night cycles with access to standard chow and water ad libitum.

## METHODS DETAILS

### Macrophage depletion

Pregnant CD1 females were given PLX5622 (Cayman Chemicals, #28927) mixed into standard chow (ScottPharma) at 1200 ppm, starting from embryonic (E) E6.5-E8.5 until E15.5-E17.5. Myeloid cell depletion was verified by immunostaining.

### Washed red blood cell preparation

Whole blood was collected from an E14.5 donor litter into an Eppendorf with 3 µL of 0.8% sodium citrate for anticoagulation. Blood was then centrifuged for 10 minutes at 500g at 4°C, and supernatant removed. This volume was measured and then 3 µL was subtracted to account for the anticoagulant volume. Approximately 1 mL of sterile saline was added to the tube and the blood resuspended before being centrifuged again. Supernatant was removed, and the wash step repeated at least once, or until supernatant was clear. Replacement of sterile saline to original volume was then added, and cells resuspended.

### Intracerebroventricular injection surgery in embryonic, postnatal, and adult animals

E14.5 pregnant dams were anesthetized. Following a laparotomy, 2 µL of fluid was delivered into one lateral ventricle per embryo. For saline injections, 0.1% fast green was added in a 1:100 dilution to visually confirm the injection was successful.^72,73^

P7 pups and adults were anesthetized and ICV delivery was achieved using pulled glass capillary needles. The following injections were performed ICV: 70 kDa TexasRed dextran (ThermoFisher, D1864, 0.25 mg/mL), 70 kDa Biotin-dextran (ThermoFisher, D1957, 5 mg/mL) + streptavidin-647 (ThermoFisher, S21374, 0.5 mg/mL), 70 kDa Biotin-dextran (5 mg/mL) + streptavidin-pacific blue (ThermoFisher, S11222, 0.5 mg/mL), or washed red blood cells (see above protocol).

70 kDa TexasRed dextran was concurrently used to mark stromal macrophages in some experiments. These injections used 25 mg/mL concentrated dextran, and were delivered intraperitoneally (P7, ∼5µL; adult, 100 µL) or intraplacentally (E14.5, 5-7µL).

### Embryonic live imaging protocol

Timed-pregnant mice (E12.5 or E15.5) were bred using female CD1 animals (CRL) and male *Cx3cr1^GFP^***^/^***^GFP^* (RRID:IMSR_JAX:005582), which allowed for the visualization of macrophages using GFP and maximized litter size. Dams were anesthetized, laparotomies were performed, and embryos were gently exposed. Vasculature labeling was achieved by placental injection of ∼7.5 μl of 25 mg/ml Texas Red-Dextran (70 kD, ThermoFisher Scientific, D1864). Following a 10 minute wait to allow the dextran to circulate, the samples with the placenta were transferred to an imaging chamber filled with artificial CSF (aCSF) at 37°C (CSHL formulation: https://cshprotocols.cshlp.org/content/2011/9/pdb.rec065730.full, 119 mM NaCl, 2.5 mM KCL, 1 mM NaH_2_PO_4_, 26.2 mM NaHCO_3_, 1.3 mM MgCl_2_, 2.5 mM CaCl_2_,10 mM glucose for E15.5). The lateral ventricle and choroid plexus (ChP) were exposed. Oxygenated (95/5 O_2_/CO_2_) and warmed aCSF was continuously circulated in the imaging chamber to maintain physiological temperature.

Two-photon imaging of immune cells was performed using either resonant-scanning (512 × 512 pixels/frame, 8.1 frames/s) or Galvo-scanning (1024x1024 pixels/frame, 1.3 frames/s) two-photon microscope (Olympus MPE-RS Multiphoton Microscope, Cellular Imaging Core, Harvard Medical School) and a 25X objective (Olympus XLSLPLN25XSVMP2, 0.95 NA, 8 mm W.D.), varying from 1x to 3x zoom. Recordings were between 5-30 minutes in duration. Volume scanning was achieved using a piezoelectric scanner (nPFocus250). Laser power at 940 nm (Mai Tai DeepSee laser, Spectra Physics) measured below the objective was 30-70mW; laser power was briefly used at 780 nm for single frames to capture the blue channel in Supplemental Figure S1C. Movies were registered using HyperStackReg (ImageJ, (Ved Sharma. (2018). ImageJ plugin HyperStackReg V5.6 (v5.6). Zenodo. https://doi.org/10.5281/zenodo.2252521)) and/or using previous code from the lab (https://github.com/LehtinenLab/Shipley2020)^24^.

### Immunohistochemistry on sections

Embryonic brains were fixed in 4% PFA at 4C for 90 minutes to overnight, depending on the gestational age. For postnatal and adult ages, animals were anesthetized prior to transcardial perfusion and post-fixed at 4°C overnight in 4% PFA. Cryoprotection was accomplished by placing the fixed brains in 30% sucrose/PBS for 1-3 days, prior to freezing in OCT. Brain blocks were stored at -80. Sections were cut at 14 µm thick onto slides at -20°C.

Slides were blocked in PBS with 0.3% Triton X (PBS-T) and 5% either goat or donkey serum at room temperature for 1 hour. Primary antibodies were diluted in blocking buffer and applied to slides at 4°C overnight. Following washes with PBS-T and PBS x 2 for 5 minutes each, secondary antibodies (1:1000 – 1:500) diluted in blocking buffer were applied to the slides for 1.5 hours at room temperature. Following washes again as above, Hoechst (ThermoFisher Scientific, H3570, 1:5000 in PBS) was applied to the slides for 10 minutes. For some slides, a drop of TrueBlack (prepared as per manufacturer’s instructions, Biotium, 23007) was applied to each section on the slide for roughly 60 seconds, before being thoroughly washed with PBS. All slides were wicked of excess liquid and mounted with Fluoromount G (ThermoFisher Scientific, OB100-01).

The following primary antibodies were used: chicken anti-GFP (Abcam ab13970; 1:1000), rabbit anti-RFP (Rockland 600-401-379, 1:500), rat anti-PECAM1 (BD Pharmingen 550274, 1:100), rat anti-CD45 (Fisher Scientific BDB550539, 1:50), rabbit anti-IBA1 (Wako 019-19741, 1:200), Goat anti-mouse MCSF antibody (R&D AF416, 1:100), Rat anti-CD68 (Abcam, ab53444, 1:250), Anti-LYVE1 conj. 488 (ThermoFisher, 53-0443-82, 1:100), Goat anti-OPN (SPP1) (Novus Biologicals, AF808, 1:100), Goat anti-GPNMB (R&D Systems, AF2330, 1:500), Rat anti-LY76 (Ter119)(Abcam, ab91113, 1:500), Chicken anti-IBA1 (Synaptic Systems, 234-009, 1:500), Rabbit anti-AQP1 (Millipore Sigma, AB2219, 1:100), Rabbit anti-MRC1 (CD206) (Abcam, ab64693, 1:250), Rabbit anti-HMOX1 (Proteintech, 10701-1-AP, 1:250).

Secondary antibodies were selected from the Alexa series (Invitrogen, 1:1000-1:500), except for the anti-goat 647 (Jackson Immunoresearch, 705-605-147, 1:500). Images were acquired using a Zeiss LSM880 or LSM980 confocal microscope.

### Quantitative PCR

Choroid plexus was pooled and RNA extracted with RNAEasy Microprep kit (Qiagen Cat# no. 74004) as per the manufacturer’s protocol. RNA was quantified with nanodrop and normalized to 100ng input and cDNA synthesis was done with Lunascript RT Supermix kit (NEB, Cat# E3010) following the manufacturer’s protocol. cDNA was amplified with 20 ul total volume of Gene Expression Mastermix (Thermo Fisher, Cat# 4369016) and 5ul of cDNA using predesigned Hmox1 (Mm00516005_m1) or Nramp1 (Mm00443045_m1) primers (Integrated DNA Technologies).

### Iron staining

Slides were washed in DI water to remove OCT prior to staining. Perl’s Prussian blue staining was made by mixing 5% potassium ferrocyanide (in DI water) combined with 5% aqueous hydrochloric acid in a 1:1 ratio. This stain was then applied to the slides for 30 minutes. Slides were then rinsed in DI water and counterstained with pararosaniline (Iron Stain Kit, Sigma-Aldrich, HT20-1KT, according to manufacturer’s instructions), prior to dehydration (1 minute each in 70%, 90%, 100% ethanol, followed by 2 minutes in xylene, twice), and mounting with Permount (Electron Microscopy Sciences, 17986-01).

For GFP signal acquisition before iron stain, P1 brains from *Cx3cr1*^GFP/+^ animals were imaged after injection of whole blood at E14.5, cryoprotection, and sectioning. Imaging was done on a Zeiss 710 (CIC Imaging Core, Boston Children’s Hospital) with no coverslip or mounting. Slides were then stained with Perls Prussian Blue as above and reimaged on a widefield light microscope (Olympus Bx51).

### Explant preparation on slides

E14.5 and P7 choroid plexus explants were dissected (E14.5 fixed for 90 min prior to dissection; P7s fresh) and postfixed in 2% PFA overnight. Explants were washed briefly in PBS and placed in Hoechst 1:5000 in PBS for 12 minutes. Explants were then rinsed in PBS and mounted on a glass slide. Extra PBS was wicked away, and the slide mounted with Fluoromount G and a coverslip. Confocal images were acquired using the Zeiss LSM 880 with Airyscan Fast (Cell Function and Imaging Core, Boston Children’s Hospital), or the Zeiss LSM710, or the Zeiss LSM 980 confocal (both of Cellular Imaging Core, Boston Children’s Hospital).

### ImageStream dissociation

Dissected and pooled LV from 1 litter of E15.5 CD1 WT embryos, previously injected 24 hours prior with 0.25 mg/mL 70 kD Texas Red dextran, were placed into digestion solution (HBSS w/no phenol red + 1 mg/mL collagenase (ThermoFisher Scientific, C6885) and 1.5 mg/mL dispase II (Gibco™ Dispase II, ThermoFisher Scientific, 17-105-041)) for 30 mins at 37° C with shaking at 600 rpm. Then, DNaseI (Worthington, LK003172) was added (4 µl for 100 µl digestion solution) and incubated for 15 min. 1 ml DMEM was added to stop enzyme activity, and cells were collected by centrifuging (10 min. at 400g, 4° C). These cells were then washed with 3% BSA in PBS and resuspended, followed by centrifuging again (10 min. at 400g, 4°C). The cells were first stained for outer markers with Iba1 (Iba1/AIF-1 (E4O4W) XP® Rabbit mAb (Alexa Fluor® 488 Conjugate) #20825) in 0.5%BSA/HBSS for 25 min. on ice, washed as above with 3% BSA and then fixed with Cytofix/Cytoperm (BD bioscience, cat # 554714, 50 µl for sample for 20 min. at room temperature). Next, cells were incubated on ice with Hoechst 1:1000 (Sigma # 23491-45-4) and washed as above one last time, then resuspended in 40µL 0.5% BSA prior to analysis with ImageStream (Flow Cytometry PCMM core facility at Boston Children’s Hospital and Harvard Medical School).

### ChP dissociation and flow cytometry sorting

Digestion buffer: 0.5 mM EDTA in HBSS, 1 mg/mL collagenase P, 1.5 mg/mL dispase II

Wash Buffer: 0.5 mM EDTA in HBSS with 3%BSA

Flow buffer: PBS/3% FBS with 0.1% EDTA

Inhibition enzymes^74^: Anisomycin (Millipore Sigma, A9789) added at 5µg/mL, actinomycin D (Millipore Sigma, A1410) added at 5 µg/mL, and triptolide (Millipore Sigma, T3652) used at 10 µM.

Digestion buffer was prepared as above with inhibition enzymes added. ChP from E14.5 and P7 animals previously injected ICV with dextran-biotin + streptavidin 647 (see above for protocol) and injected IP with 70 kD Texas Red dextran were collected into 500uL of buffer. One litter of paired lateral ChP were pooled. These were placed on a rotator at 37°C for digestion for 15 minutes, with gentle pipetting to help dissociation every 5 minutes. DNase I was added after 15 minutes. After another 15 minutes, still pipetting gently every 5, 1 mL RPMI was added. Collection vial was then centrifuged at 4C at 100g for 10 minutes. Supernatant was removed from the tube and replaced with 300 µL of wash buffer and pellet resuspended with tips coated in PBS/3%BSA. Collection tube was centrifuged again as above and washed a second time. After final spin, supernatant was replaced with flow buffer and placed on ice. Cell sorting was done on a BD FACSARIA III (Flow Cytometry Core, Boston Children’s Hospital) directly into RLT buffer. RNA was extracted by proceeding with the RNeasy Micro kit (Qiagen, 74004), according to manufacturer’s instructions.

### Library generation, sequencing, and analysis

Libraries for macrophages were prepared using the CORALL mRNA-Seq V2 Library Prep Kit with UDI 12 nt Set B1 (cat# 181.96, Lexogen, Vienna, Austria) following the manufacturer’s protocol. Sequencing was on a NovaSeq S1 200 cycle run. Unique molecular identifiers were used to deduplicate reads using umi_tools 1.0.0 (https://github.com/CGATOxford/UMI-tools) and trimmed using cutadapt v2.5.^75^ Processed reads were aligned to the Ensembl mouse genome GRMCm39 using STAR 2.6.1b^76^ and quantified with featureCounts v.2.0.6.^77^ Samtools 1.9 was used for indexing and formatting.^78^ FastQC v0.11.8 (https://www.bioinformatics.babraham.ac.uk/projects/fastqc/) and multiQC v1.7^79^ were used at every processing step for quality control. Statistical analysis was done in R version 4.3.3 using DESeq2 v1.42.1.^80^ Dimension reduction was done using variance stabilizing transformation and plotted using ggplot2.^81^ Gene ontology was done using ClusterProfiler v4.10.1.^82^ Choroid plexus samples were sent to Novogene for sequencing after extraction.

### Bulk RNA-seq analyses

Gene set and Reactome enrichment were performed using ClusterProfiler v4.10.1.^82^ To infer cell type contributions to bulk transcriptional profile, we applied the MuSiC algorithm^83^ using the previously published scRNA-seq data from Xu et al.^31^ Cell clusters were aggregated into biologically meaningful categories (e.g., epithelial subtypes, fibroblasts, endothelial, macrophages, neutrophils, dendritic cells, cytotoxic effector cells, and monocytes) prior to deconvolution. Weighted cell type proportions were estimated for each bulk sample and compared across conditions. Visualization was performed using ggplot2, including bar plots stratified by condition. To further delineate the cell-types driving DEG, we used a principle component-based analysis projected onto single cell data following the previously reported STAR protocol.^84^

### Analysis of scRNA-Seq

Single-cell RNA-seq data from Dani and Herbst et al. 2021 were processed using Seurat (v4).^85^ Raw count matrices and associated gene and cell barcode annotations were imported, and mitochondrial RNA content was calculated to filter low-quality cells. Cells from replicate groups E.3V.V2.Rep2, E.HCP.V2.Rep2, and E.TCP.V2.Rep2, as well as cells prepared using the EFO_0009901 library protocol, were excluded due to high mitochondrial RNA content. Macrophages were filtered based on previous annotations and normalized using log-normalization. Highly variable genes were identified (n = 2,000), and data were scaled prior to principal component (PC) analysis. Significant PCs were selected for clustering and dimensionality reduction using UMAP. Cluster-specific marker genes were identified using a log-fold change threshold of 1 and minimum expression in 25% of cells. Cell markers were chosen based on known tissue-and origin-specific relevance (e.g., *Spic+Hmox1, Lyve1+Cd163, Cd9+Spp1+Gpnmb, Nusap+Ki67*), and visualizations were generated using DimPlot() in Seurat and pheatmap.^86^ Gene ontology designation for each cell cluster was done using https://www.geneontology.org/.^87,88^ Ligand–receptor signaling networks among choroid plexus cell populations were inferred using CellChat.^37,38^

To identify common signatures with tissue-and fluid-niche macrophages we defined gene set for each population based on previous literature (LPM^39,41^: *Gata6, Timd4, Mertk, Klf2, Cd93, Vsig4, Marco, Cd1c, Selp*; SPM^39,41^: *Cd226, Irf4, Il1b, Tnf, H2-Ab1, Ccr2, Lyve1, Mrc1, Csf1r*; sentinel^89,90^: *Itgam, Fcgr1, Cd68, Cd80, Cd86, Tlr2, Tlr4, Msr1*; BAM^91^: *Pf4, F13a1, Cd163, Flt1, Lyve1, Mrc1*; microglia^92^: *P2ry12, Tmem119, Sall1, Siglech, Hexb, Fcrls, Gpr34, Cx3cr1, Axl, Crybb1, Ccl4*; monocyte: *Cd14, Lyz, Fcn1, S100a8, S100a9*; YAMs^44^: *Spp1, Gpnmb, Cst7, Fabp5, Lgals3, Apoe, Trem2, Clec7a*). Module scores summarizing the average expression of each gene set were calculated per cell to quantify cell-type specific transcriptional programs using Seurat v4 with appropriate plotting features, and differences across clusters were assessed using Kruskal-Wallis.^93^

## QUANTIFICATION AND STATISTICAL ANALYSIS

### Statistics and Quantification

Biological replicates (N) were samples from distinct embryos. Sample sizes were informed by estimated mean values from preliminary data and previous studies. Cell counts were performed in a blinded manner, and all cell counts were averaged from between 3-12 lateral ventricle slices per animal, depending on the experiment. Percent of ventricle occupied by RBCs in PLX vs control experiment was done by measuring area of the ventricle, subtracting the ChP area, and then thresholding the RBC signal within the ventricle ROI for rough area. Two slices were used to average per animal. Statistical analyses were performed using Prism 10 (GraphPad). Statistical significance between two groups was done using Welch’s unpaired *t*-test except for the epiplexus/intraventricular count comparisons across ages, which was evaluated using a Kruskal-Wallis ANOVA with Dunnett’s multiple comparisons test. Data are presented as mean ± standard deviation (SD). Each datapoint represents one biological replicate and was plotted separately in figures. All datapoints were included (no outliers removed). See Results or figure legends for statistical tests and figure panels for sample sizes for each experiment. p values < 0.05 were considered significant (*p <0.05, ** p <0.01, *** p <0.001).

## REFERENCES

1. Courtney, Y., Head, J.P., Dani, N., Chechneva, O.V., Shipley, F.B., Zhang, Y., Holtzman, M.J., Sadegh, C., Libermann, T.A., and Lehtinen, M.K. (2025). Choroid plexus apocrine secretion shapes CSF proteome during mouse brain development. Nat Neurosci 28, 1446– 1459. 10.1038/s41593-025-01972-9.

2. Gelb, S., and Lehtinen, M.K. (2023). Snapshot: Choroid plexus brain barrier. Cell 186, 3522–3522.e1. 10.1016/j.cell.2023.07.015.

3. Lazarevic, I., Soldati, S., Mapunda, J.A., Rudolph, H., Rosito, M., de Oliveira, A.C., Enzmann, G., Nishihara, H., Ishikawa, H., Tenenbaum, T., et al. (2023). The choroid plexus acts as an immune cell reservoir and brain entry site in experimental autoimmune encephalomyelitis. Fluids Barriers CNS 20, 39. 10.1186/s12987-023-00441-4.

4. Xu, H., Hehnly, C., and Lehtinen, M.K. (2025). The choroid plexus: a command center for brain–body communication during inflammation. Current Opinion in Immunology 93, 102540. 10.1016/j.coi.2025.102540.

5. De Vlaminck, K., Van Hove, H., Kancheva, D., Scheyltjens, I., Pombo Antunes, A.R., Bastos, J., Vara-Perez, M., Ali, L., Mampay, M., Deneyer, L., et al. (2022). Differential plasticity and fate of brain-resident and recruited macrophages during the onset and resolution of neuroinflammation. Immunity 55, 2085–2102.e9. 10.1016/j.immuni.2022.09.005.

6. Munro, D.A.D., Movahedi, K., and Priller, J. (2022). Macrophage compartmentalization in the brain and cerebrospinal fluid system. Sci Immunol 7, eabk0391. 10.1126/sciimmunol.abk0391.

7. Leijser, L.M., and De Vries, L.S. (2019). Preterm brain injury: Germinal matrix– intraventricular hemorrhage and post-hemorrhagic ventricular dilatation. In Handbook of Clinical Neurology (Elsevier), pp. 173–199. 10.1016/B978-0-444-64029-1.00008-4.

8. Ballabh, P., and de Vries, L.S. (2021). White matter injury in infants with intraventricular haemorrhage: mechanisms and therapies. Nat Rev Neurol 17, 199–214. 10.1038/s41582-020-00447-8.

9. Rees, P., Gale, C., Battersby, C., Williams, C., Carter, B., and Sutcliffe, A. (2025). Intraventricular Hemorrhage and Survival, Multimorbidity, and Neurodevelopment. JAMA Network Open 8, e2452883. 10.1001/jamanetworkopen.2024.52883.

10. Nagy, Z., Obeidat, M., Máté, V., Nagy, R., Szántó, E., Veres, D.S., Kói, T., Hegyi, P., Major, G.S., Garami, M., et al. (2025). Occurrence and Time of Onset of Intraventricular Hemorrhage in Preterm Neonates: A Systematic Review and Meta-Analysis of Individual Patient Data. JAMA Pediatr 179, 145. 10.1001/jamapediatrics.2024.5998.

11. Juliet, P.A.R., Frost, E.E., Balasubramaniam, J., and Del Bigio, M.R. (2009). Toxic effect of blood components on perinatal rat subventricular zone cells and oligodendrocyte precursor cell proliferation, differentiation and migration in culture. Journal of Neurochemistry 109, 1285–1299. 10.1111/j.1471-4159.2009.06060.x.

12. Gram, M., Sveinsdottir, S., Cinthio, M., Sveinsdottir, K., Hansson, S.R., Mörgelin, M., Åkerström, B., and Ley, D. (2014). Extracellular hemoglobin - mediator of inflammation and cell death in the choroid plexus following preterm intraventricular hemorrhage. J Neuroinflammation 11, 200. 10.1186/s12974-014-0200-9.

13. Ye, F., Hua, Y., Keep, R.F., Xi, G., and Garton, H.J.L. (2021). CD47 blocking antibody accelerates hematoma clearance and alleviates hydrocephalus after experimental intraventricular hemorrhage. Neurobiol Dis 155, 105384. 10.1016/j.nbd.2021.105384.

14. Yauger, Y.J., Bermudez, S., Moritz, K.E., Glaser, E., Stoica, B., and Byrnes, K.R. (2019). Iron accentuated reactive oxygen species release by NADPH oxidase in activated microglia contributes to oxidative stress in vitro. J Neuroinflammation 16, 41. 10.1186/s12974-019-1430-7.

15. Mahaney, K.B., Buddhala, C., Paturu, M., Morales, D., Limbrick, D.D., and Strahle, J.M. (2020). Intraventricular Hemorrhage Clearance in Human Neonatal Cerebrospinal Fluid: Associations with Hydrocephalus. Stroke 51, 1712–1719. 10.1161/STROKEAHA.119.028744.

16. Strahle, J.M., Mahaney, K.B., Morales, D.M., Buddhala, C., Shannon, C.N., Wellons III, J.C., Kulkarni, A.V., Jensen, H., Reeder, R.W., Holubkov, R., et al. (2021). Longitudinal CSF Iron Pathway Proteins in Posthemorrhagic Hydrocephalus: Associations with Ventricle Size and Neurodevelopmental Outcomes. Annals of Neurology 90, 217–226. 10.1002/ana.26133.

17. Strahle, J.M., Garton, T., Bazzi, A.A., Kilaru, H., Garton, H.J.L., Maher, C.O., Muraszko, K.M., Keep, R.F., and Xi, G. (2014). Role of hemoglobin and iron in hydrocephalus after neonatal intraventricular hemorrhage. Neurosurgery 75, 696–705; discussion 706. 10.1227/NEU.0000000000000524.

18. Mrdjen, D., Pavlovic, A., Hartmann, F.J., Schreiner, B., Utz, S.G., Leung, B.P., Lelios, I., Heppner, F.L., Kipnis, J., Merkler, D., et al. (2018). High-Dimensional Single-Cell Mapping of Central Nervous System Immune Cells Reveals Distinct Myeloid Subsets in Health, Aging, and Disease. Immunity 48, 380–395.e6. 10.1016/j.immuni.2018.01.011.

19. Kolmer, W. (1921). Über eine eigenartige Beziehung von Wanderzellen zu den Chorioidealplexus des Gehirns der Wirbeltiere. Anat. Anz 54, 15–19.

20. Ling, E.-A., Kaur, C., and Lu, J. (1998). Origin, nature, and some functional considerations of intraventricular macrophages, with special reference to the epiplexus cells. Microscopy Research and Technique 41, 43–56. 10.1002/(SICI)1097-0029(19980401)41:1%253C43::AID-JEMT5%253E3.0.CO;2-V.

21. Munro, D.A.D., Bradford, B.M., Mariani, S.A., Hampton, D.W., Vink, C.S., Chandran, S., Hume, D.A., Pridans, C., and Priller, J. (2020). CNS macrophages differentially rely on an intronic Csf1r enhancer for their development. Development 147, dev194449. 10.1242/dev.194449.

22. Van Hove, H., Martens, L., Scheyltjens, I., De Vlaminck, K., Pombo Antunes, A.R., De Prijck, S., Vandamme, N., De Schepper, S., Van Isterdael, G., Scott, C.L., et al. (2019). A single-cell atlas of mouse brain macrophages reveals unique transcriptional identities shaped by ontogeny and tissue environment. Nat Neurosci 22, 1021–1035. 10.1038/s41593-019-0393-4.

23. McKinsey, G.L., Lizama, C.O., Keown-Lang, A.E., Niu, A., Santander, N., Larpthaveesarp, A., Chee, E., Gonzalez, F.F., and Arnold, T.D. (2020). A new genetic strategy for targeting microglia in development and disease. eLife 9, e54590. 10.7554/eLife.54590.

24. Shipley, F.B., Dani, N., Xu, H., Deister, C., Cui, J., Head, J.P., Sadegh, C., Fame, R.M., Shannon, M.L., Flores, V.I., et al. (2020). Tracking Calcium Dynamics and Immune Surveillance at the Choroid Plexus Blood-Cerebrospinal Fluid Interface. Neuron 108, 623–639.e10. 10.1016/j.neuron.2020.08.024.

25. Cui, J., Shipley, F.B., Shannon, M.L., Alturkistani, O., Dani, N., Webb, M.D., Sugden, A.U., Andermann, M.L., and Lehtinen, M.K. (2020). Inflammation of the Embryonic Choroid Plexus Barrier following Maternal Immune Activation. Dev Cell 55, 617–628.e6. 10.1016/j.devcel.2020.09.020.

26. Chen, J., Crouch, E.E., Zawadzki, M.E., Jacobs, K.A., Mayo, L.N., Choi, J.J.-Y., Lin, P.-Y., Shaikh, S., Tsui, J., Gonzalez-Granero, S., et al. (2024). Proinflammatory immune cells disrupt angiogenesis and promote germinal matrix hemorrhage in prenatal human brain. Nat Neurosci 27, 2115–2129. 10.1038/s41593-024-01769-2.

27. Davalos, D., Grutzendler, J., Yang, G., Kim, J.V., Zuo, Y., Jung, S., Littman, D.R., Dustin, M.L., and Gan, W.-B. (2005). ATP mediates rapid microglial response to local brain injury in vivo. Nat Neurosci 8, 752–758. 10.1038/nn1472.

28. Nimmerjahn, A., Kirchhoff, F., and Helmchen, F. (2005). Resting Microglial Cells Are Highly Dynamic Surveillants of Brain Parenchyma in Vivo. Science 308, 1314–1318. 10.1126/science.1110647.

29. Milacic, M., Beavers, D., Conley, P., Gong, C., Gillespie, M., Griss, J., Haw, R., Jassal, B., Matthews, L., May, B., et al. (2024). The Reactome Pathway Knowledgebase 2024. Nucleic Acids Res 52, D672–D678. 10.1093/nar/gkad1025.

30. Dani, N., Herbst, R.H., McCabe, C., Green, G.S., Kaiser, K., Head, J.P., Cui, J., Shipley, F.B., Jang, A., Dionne, D., et al. (2021). A cellular and spatial map of the choroid plexus across brain ventricles and ages. Cell 184, 3056–3074.e21. 10.1016/j.cell.2021.04.003.

31. Xu, H., Lotfy, P., Gelb, S., Pragana, A., Hehnly, C., Byer, L.I.J., Shipley, F.B., Zawadzki, M.E., Cui, J., Deng, L., et al. (2024). The choroid plexus synergizes with immune cells during neuroinflammation. Cell, S0092867424007177. 10.1016/j.cell.2024.07.002.

32. Soe-Lin, S., Apte, S.S., Andriopoulos, B., Andrews, M.C., Schranzhofer, M., Kahawita, T., Garcia-Santos, D., and Ponka, P. (2009). Nramp1 promotes efficient macrophage recycling of iron following erythrophagocytosis in vivo. Proc Natl Acad Sci U S A 106, 5960–5965. 10.1073/pnas.0900808106.

33. Recalcati, S., and Cairo, G. (2021). Macrophages and Iron: A Special Relationship. Biomedicines 9, 1585. 10.3390/biomedicines9111585.

34. Wan, Y., Fu, X., Zhang, T., Hua, Y., Keep, R.F., and Xi, G. (2024). Choroid plexus immune cell response in murine hydrocephalus induced by intraventricular hemorrhage. Fluids Barriers CNS 21, 37. 10.1186/s12987-024-00538-4.

35. Elmore, M.R.P., Najafi, A.R., Koike, M.A., Dagher, N.N., Spangenberg, E.E., Rice, R.A., Kitazawa, M., Matusow, B., Nguyen, H., West, B.L., et al. (2014). CSF1 receptor signaling is necessary for microglia viability, which unmasks a cell that rapidly repopulates the microglia-depleted adult brain. Neuron 82, 380–397. 10.1016/j.neuron.2014.02.040.

36. Rosin, J.M., Vora, S.R., and Kurrasch, D.M. (2018). Depletion of embryonic microglia using the CSF1R inhibitor PLX5622 has adverse sex-specific effects on mice, including accelerated weight gain, hyperactivity and anxiolytic-like behaviour. Brain, Behavior, and Immunity 73, 682–697. 10.1016/j.bbi.2018.07.023.

37. Jin, S., Guerrero-Juarez, C.F., Zhang, L., Chang, I., Ramos, R., Kuan, C.-H., Myung, P., Plikus, M.V., and Nie, Q. (2021). Inference and analysis of cell-cell communication using CellChat. Nat Commun 12, 1088. 10.1038/s41467-021-21246-9.

38. Jin, S., Plikus, M.V., and Nie, Q. (2025). CellChat for systematic analysis of cell–cell communication from single-cell transcriptomics. Nat Protoc 20, 180–219. 10.1038/s41596-024-01045-4.

39. Cassado, A. dos A., D’Império Lima, M.R., and Bortoluci, K.R. (2015). Revisiting Mouse Peritoneal Macrophages: Heterogeneity, Development, and Function. Front Immunol 6, 225. 10.3389/fimmu.2015.00225.

40. Utz, S.G., See, P., Mildenberger, W., Thion, M.S., Silvin, A., Lutz, M., Ingelfinger, F., Rayan, N.A., Lelios, I., Buttgereit, A., et al. (2020). Early Fate Defines Microglia and Non-parenchymal Brain Macrophage Development. Cell 181, 557–573.e18. 10.1016/j.cell.2020.03.021.

41. Han, J., Gallerand, A., Erlich, E.C., Helmink, B.A., Mair, I., Li, X., Eckhouse, S.R., Dimou, F.M., Shakhsheer, B.A., Phelps, H.M., et al. (2024). Human serous cavity macrophages and dendritic cells possess counterparts in the mouse with a distinct distribution between species. Nat Immunol 25, 155–165. 10.1038/s41590-023-01688-7.

42. Hammond, T.R., Dufort, C., Dissing-Olesen, L., Giera, S., Young, A., Wysoker, A., Walker, A.J., Gergits, F., Segel, M., Nemesh, J., et al. (2019). Single cell RNA sequencing of microglia throughout the mouse lifespan and in the injured brain reveals complex cell-state changes. Immunity 50, 253–271.e6. 10.1016/j.immuni.2018.11.004.

43. Li, Q., Cheng, Z., Zhou, L., Darmanis, S., Neff, N.F., Okamoto, J., Gulati, G., Bennett, M.L., Sun, L.O., Clarke, L.E., et al. (2019). Developmental heterogeneity of microglia and brain myeloid cells revealed by deep single-cell RNA sequencing. Neuron 101, 207–223.e10. 10.1016/j.neuron.2018.12.006.

44. Silvin, A., Uderhardt, S., Piot, C., Mesquita, S.D., Yang, K., Geirsdottir, L., Mulder, K., Eyal, D., Liu, Z., Bridlance, C., et al. (2022). Dual ontogeny of disease-associated microglia and disease inflammatory macrophages in aging and neurodegeneration. Immunity 55, 1448–1465.e6. 10.1016/j.immuni.2022.07.004.

45. Hattori, Y., Kato, D., Murayama, F., Koike, S., Asai, H., Yamasaki, A., Naito, Y., Kawaguchi, A., Konishi, H., Prinz, M., et al. (2023). CD206+ macrophages transventricularly infiltrate the early embryonic cerebral wall to differentiate into microglia. Cell Reports 42. 10.1016/j.celrep.2023.112092.

46. Tseng, C.Y., Ling, E.A., and Wong, W.C. (1983). Scanning electron microscopy of amoeboid microglial cells in the transient cavum septum pellucidum in pre-and postnatal rats. J Anat 136, 251–263.

47. Lawrence, A.R., Canzi, A., Bridlance, C., Olivié, N., Lansonneur, C., Catale, C., Pizzamiglio, L., Kloeckner, B., Silvin, A., Munro, D.A.D., et al. (2024). Microglia maintain structural integrity during fetal brain morphogenesis. Cell 187, 962–980.e19. 10.1016/j.cell.2024.01.012.

48. Esaulova, E., Cantoni, C., Shchukina, I., Zaitsev, K., Bucelli, R.C., Wu, G.F., Artyomov, M.N., Cross, A.H., and Edelson, B.T. (2020). Single-cell RNA-seq analysis of human CSF microglia and myeloid cells in neuroinflammation. Neurol Neuroimmunol Neuroinflamm 7, e732. 10.1212/NXI.0000000000000732.

49. Mukherjee, G., Waris, R., Rechler, W., Kudelka, M., McCracken, C., Kirpalani, A., and Hames, N. (2021). Determining Normative Values for Cerebrospinal Fluid Profiles in Infants. Hospital Pediatrics 11, 930–936. 10.1542/hpeds.2020-005512.

50. Martín-Ancel, A., García-Alix, A., Salas, S., del Castillo, F., Cabañas, F., and Quero, J. (2006). Cerebrospinal fluid leucocyte counts in healthy neonates. Arch Dis Child Fetal Neonatal Ed 91, F357–F358. 10.1136/adc.2005.082826.

51. Verney, C., Monier, A., Fallet-Bianco, C., and Gressens, P. (2010). Early microglial colonization of the human forebrain and possible involvement in periventricular white-matter injury of preterm infants. J Anat 217, 436–448. 10.1111/j.1469-7580.2010.01245.x.

52. Kracht, L., Borggrewe, M., Eskandar, S., Brouwer, N., Chuva de Sousa Lopes, S.M., Laman, J.D., Scherjon, S.A., Prins, J.R., Kooistra, S.M., and Eggen, B.J.L. (2020). Human fetal microglia acquire homeostatic immune-sensing properties early in development. Science 369, 530–537. 10.1126/science.aba5906.

53. Keren-Shaul, H., Spinrad, A., Weiner, A., Matcovitch-Natan, O., Dvir-Szternfeld, R., Ulland, T.K., David, E., Baruch, K., Lara-Astaiso, D., Toth, B., et al. (2017). A Unique Microglia Type Associated with Restricting Development of Alzheimer’s Disease. Cell 169, 1276–1290.e17. 10.1016/j.cell.2017.05.018.

54. Brioschi, S., Zhou, Y., and Colonna, M. (2020). Brain Parenchymal and Extraparenchymal Macrophages in Development, Homeostasis, and Disease. J Immunol 204, 294–305. 10.4049/jimmunol.1900821.

55. Martins-Ferreira, R., Calafell-Segura, J., Leal, B., Rodríguez-Ubreva, J., Martínez-Saez, E., Mereu, E., Pinho E Costa, P., Laguna, A., and Ballestar, E. (2025). The Human Microglia Atlas (HuMicA) unravels changes in disease-associated microglia subsets across neurodegenerative conditions. Nat Commun 16, 739. 10.1038/s41467-025-56124-1.

56. Hohsfield, L.A., Kim, S.J., Barahona, R.A., Henningfield, C.M., Mansour, K., Vallejo, K.D., Tsourmas, K.I., Kwang, N.E., Ghorbanian, Y., Angulo, J.A.A., et al. (2025). Identification of the velum interpositum as a meningeal-CNS route for myeloid cell trafficking into the brain. Neuron 113, 2455–2473.e6. 10.1016/j.neuron.2025.05.004.

57. Wang, J., and Kubes, P. (2016). A Reservoir of Mature Cavity Macrophages that Can Rapidly Invade Visceral Organs to Affect Tissue Repair. Cell 165, 668–678. 10.1016/j.cell.2016.03.009.

58. Whitelaw, A., Jary, S., Kmita, G., Wroblewska, J., Musialik-Swietlinska, E., Mandera, M., Hunt, L., Carter, M., and Pople, I. (2010). Randomized Trial of Drainage, Irrigation and Fibrinolytic Therapy for Premature Infants with Posthemorrhagic Ventricular Dilatation: Developmental Outcome at 2 years. Pediatrics 125, e852–e858. 10.1542/peds.2009-1960.

59. Etus, V., Kahilogullari, G., Karabagli, H., and Unlu, A. (2016). Early endoscopic ventricular irrigation for the treatment of neonatal posthemorrhagic hydrocephalus. a feasible treatment option or not□? - a multi center report-. Turkish Neurosurgery. 10.5137/1019-5149.JTN.18677-16.0.

60. Luyt, K., Jary, S.L., Lea, C.L., Young, G.J., Odd, D.E., Miller, H.E., Kmita, G., Williams, C., Blair, P.S., Hollingworth, W., et al. (2020). Drainage, irrigation and fibrinolytic therapy (DRIFT) for posthaemorrhagic ventricular dilatation: 10-year follow-up of a randomised controlled trial. Arch Dis Child Fetal Neonatal Ed 105, 466–473. 10.1136/archdischild-2019-318231.

61. Chen, J., Wang, L., Xu, H., Xing, L., Zhuang, Z., Zheng, Y., Li, X., Wang, C., Chen, S., Guo, Z., et al. (2020). Meningeal lymphatics clear erythrocytes that arise from subarachnoid hemorrhage. Nat Commun 11, 3159. 10.1038/s41467-020-16851-z.

62. Bocheng, W., Chaofeng, L., Chuan, C., Haiyong, H., Tengchao, H., Qun, G., and Ying, G. (2020). Intracranial lymphatic drainage discharges iron from the ventricles and reduce the occurrence of chronic hydrocephalus after intraventricular hemorrhage in rats. International Journal of Neuroscience 130, 130–135. 10.1080/00207454.2019.1667780.

63. Antila, S., Karaman, S., Nurmi, H., Airavaara, M., Voutilainen, M.H., Mathivet, T., Chilov, D., Li, Z., Koppinen, T., Park, J.-H., et al. (2017). Development and plasticity of meningeal lymphatic vessels. J Exp Med 214, 3645–3667. 10.1084/jem.20170391.

64. Ma, Q., Ineichen, B.V., Detmar, M., and Proulx, S.T. (2017). Outflow of cerebrospinal fluid is predominantly through lymphatic vessels and is reduced in aged mice. Nat Commun 8, 1434. 10.1038/s41467-017-01484-6.

65. Garton, T., Keep, R.F., Hua, Y., and Xi, G. (2017). CD163, a Hemoglobin/Haptoglobin Scavenger Receptor, After Intracerebral Hemorrhage: Functions in Microglia/Macrophages Versus Neurons. Transl Stroke Res 8, 612–616. 10.1007/s12975-017-0535-5.

66. Chang, C.-F., Massey, J., Osherov, A., da Costa, L.H.A., and Sansing, L.H. (2020). Bexarotene enhances macrophage erythrophagocytosis and hematoma clearance in experimental intracerebral hemorrhage. Stroke 51, 612–618. 10.1161/STROKEAHA.119.027037.

67. Rouault, T.A., Zhang, D.-L., and Jeong, S.Y. (2009). Brain iron homeostasis, the choroid plexus, and localization of iron transport proteins. Metab Brain Dis 24, 673–684. 10.1007/s11011-009-9169-y.

68. Ramagiri, S., Pan, S., DeFreitas, D., Yang, P.H., Raval, D.K., Wozniak, D.F., Esakky, P., and Strahle, J.M. (2023). Deferoxamine Prevents Neonatal Posthemorrhagic Hydrocephalus Through Choroid Plexus-Mediated Iron Clearance. Transl. Stroke Res. 14, 704–722. 10.1007/s12975-022-01092-7.

69. Robert, S.M., Reeves, B.C., Kiziltug, E., Duy, P.Q., Karimy, J.K., Mansuri, M.S., Marlier, A., Allington, G., Greenberg, A.B.W., DeSpenza, T., et al. (2023). The choroid plexus links innate immunity to CSF dysregulation in hydrocephalus. Cell 186, 764–785.e21. 10.1016/j.cell.2023.01.017.

70. Wan, Y., Hua, Y., Garton, H.J.L., Novakovic, N., Keep, R.F., and Xi, G. (2019). Activation of epiplexus macrophages in hydrocephalus caused by subarachnoid hemorrhage and thrombin. CNS Neurosci Ther 25, 1134–1141. 10.1111/cns.13203.

71. Evangelista, J.E., Xie, Z., Marino, G.B., Nguyen, N., Clarke, D.J.B., and Ma’ayan, A. (2023). Enrichr-KG: bridging enrichment analysis across multiple libraries. Nucleic Acids Res 51, W168–W179. 10.1093/nar/gkad393.

72. Jang, A., and Lehtinen, M.K. (2022). Experimental approaches for manipulating choroid plexus epithelial cells. Fluids Barriers CNS 19, 36. 10.1186/s12987-022-00330-2.

73. Sadegh, C., Xu, H., Sutin, J., Fatou, B., Gupta, S., Pragana, A., Taylor, M., Kalugin, P.N., Zawadzki, M.E., Alturkistani, O., et al. (2023). Choroid plexus-targeted NKCC1 overexpression to treat post-hemorrhagic hydrocephalus. Neuron 111, 1591–1608.e4. 10.1016/j.neuron.2023.02.020.

74. Marsh, S.E., Walker, A.J., Kamath, T., Dissing-Olesen, L., Hammond, T.R., de Soysa, T.Y., Young, A.M.H., Murphy, S., Abdulraouf, A., Nadaf, N., et al. (2022). Dissection of artifactual and confounding glial signatures by single-cell sequencing of mouse and human brain. Nat Neurosci 25, 306–316. 10.1038/s41593-022-01022-8.

75. Martin, Marcel (2011). Cutadapt removes adapter sequences from high-throughput sequencing reads. EMBnet.jornal 17, 10–12.

76. Dobin, A., Davis, C.A., Schlesinger, F., Drenkow, J., Zaleski, C., Jha, S., Batut, P., Chaisson, M., and Gingeras, T.R. (2013). STAR: ultrafast universal RNA-seq aligner. Bioinformatics 29, 15–21. 10.1093/bioinformatics/bts635.

77. Liao, Y., Smyth, G.K., and Shi, W. (2014). featureCounts: an efficient general purpose program for assigning sequence reads to genomic features. Bioinformatics 30, 923–930. 10.1093/bioinformatics/btt656.

78. Danecek, P., Bonfield, J.K., Liddle, J., Marshall, J., Ohan, V., Pollard, M.O., Whitwham, A., Keane, T., McCarthy, S.A., Davies, R.M., et al. (2021). Twelve years of SAMtools and BCFtools. GigaScience 10, giab008. 10.1093/gigascience/giab008.

79. Ewels, P., Magnusson, M., Lundin, S., and Käller, M. (2016). MultiQC: summarize analysis results for multiple tools and samples in a single report. Bioinformatics 32, 3047–3048. 10.1093/bioinformatics/btw354.

80. Love, M.I., Huber, W., and Anders, S. (2014). Moderated estimation of fold change and dispersion for RNA-seq data with DESeq2. Genome Biology 15, 550. 10.1186/s13059-014-0550-8.

81. Wickham, H. (2016). ggplot2 (Springer International Publishing) 10.1007/978-3-319-24277-4.

82. Wu, T., Hu, E., Xu, S., Chen, M., Guo, P., Dai, Z., Feng, T., Zhou, L., Tang, W., Zhan, L., et al. (2021). clusterProfiler 4.0: A universal enrichment tool for interpreting omics data. Innovation 2. 10.1016/j.xinn.2021.100141.

83. Wang, X., Park, J., Susztak, K., Zhang, N.R., and Li, M. (2019). Bulk tissue cell type deconvolution with multi-subject single-cell expression reference. Nat Commun 10, 380. 10.1038/s41467-018-08023-x.

84. Marquez-Galera, A., De La Prida, L.M., and Lopez-Atalaya, J.P. (2022). A protocol to extract cell-type-specific signatures from differentially expressed genes in bulk-tissue RNA-seq. STAR Protocols 3, 101121. 10.1016/j.xpro.2022.101121.

85. Hao, Y., Hao, S., Andersen-Nissen, E., Mauck, W.M., Zheng, S., Butler, A., Lee, M.J., Wilk, A.J., Darby, C., Zager, M., et al. (2021). Integrated analysis of multimodal single-cell data. Cell 184, 3573–3587.e29. 10.1016/j.cell.2021.04.048.

86. Kolde, R. (2025). pheatmap: Pretty Heatmaps.

87. Ashburner, M., Ball, C.A., Blake, J.A., Botstein, D., Butler, H., Cherry, J.M., Davis, A.P., Dolinski, K., Dwight, S.S., Eppig, J.T., et al. (2000). Gene Ontology: tool for the unification of biology. Nat Genet 25, 25–29. 10.1038/75556.

88. Aleksander, S.A., Balhoff, J., Carbon, S., Cherry, J.M., Drabkin, H.J., Ebert, D., Feuermann, M., Gaudet, P., Harris, N.L., Hill, D.P., et al. (2023). The Gene Ontology knowledgebase in 2023. Genetics 224, iyad031. 10.1093/genetics/iyad031.

89. Fritz, J.H., Ferrero, R.L., Philpott, D.J., and Girardin, S.E. (2006). Nod-like proteins in immunity, inflammation and disease. Nat Immunol 7, 1250–1257. 10.1038/ni1412.

90. Palm, N.W., and Medzhitov, R. (2009). Pattern recognition receptors and control of adaptive immunity. Immunological Reviews 227, 221–233. 10.1111/j.1600-065X.2008.00731.x.

91. Sun, R., and Jiang, H. (2024). Border-associated macrophages in the central nervous system. Journal of Neuroinflammation 21, 67. 10.1186/s12974-024-03059-x.

92. Jurga, A.M., Paleczna, M., and Kuter, K.Z. (2020). Overview of General and Discriminating Markers of Differential Microglia Phenotypes. Front. Cell. Neurosci. 14. 10.3389/fncel.2020.00198.

93. Kruskal, W.H., and Wallis, W.A. (1952). Use of Ranks in One-Criterion Variance Analysis. Journal of the American Statistical Association 47, 583–621. 10.2307/2280779.

