## Supplemental text for "Fluid-Niche and Microglial Signatures Prime Robust Intraventricular Macrophage Response to Blood During Brain Development"

**SUPPLEMENTAL MATERIAL**

**Supplemental Video S1. Top-down view of E15.5 ChP in the ventricle *ex vivo* in an E15.5 *Cx3cr1*^GFP/+^ embryo, related to Figure 1L**. Still image and legend in Figure 1N. Vessels = magenta; CX3CR1(GFP) = green. Scale bar = 200 µm.

**Supplemental Video S2. Comparison of ChP macrophages and microglial movements in E15.5 *Cx3cr1*^GFP/+^ embryo from Figure 1L**. Vessels = magenta; GFP = green. Scale bar = 25 µm.

**Supplemental Video S3. Stromal vs epiplexus cell movement examples in E15.5 *Cx3cr1*^GFP/+^ mouse embryo ChP, related to Figure 1N.** *Left***:** Video magnification from **Figure 1N**; left video shows the vessels (in magenta) and macrophages (in green) and identifies example stromal cells (red arrows) and epiplexus cells (blue arrows). Right video shows only the macrophage signal (GFP = gray). Scale bar = 50 µm. *Right:* Stills represented in **Figure 1N**. Scale bar = 25 µm.

**Supplemental Video S4. ChP macrophages in an E15.5 *Cx3cr1*^GFP/+^ mouse embryo injected with saline vs RBCs 24 hours prior, related to Figure 2J.** Scale bar = 50 µm.

**Supplemental Figure S1**. (**A-B**) Dual-dextran injected ChP whole mount at E14.5, 3 hours post-injection (A) and single dextran injection IP ChP whole mount in adult, 24 hours later (B). (**C**) Still image from 2-photon video in E14.5 embryo, dual-injected with IP (red) and ICV (blue) dextrans 3 hours prior to imaging. Red arrows = stromal cells, white arrows = epiplexus cells. (**D**) HMOX1 staining in lateral ventricle 24 hours after ICV saline injection into E14.5 embryo. Solid line = ventricle, dashed line = ChP. Contrast adjusted for visualization. (**E**) *Slc11a1* qPCR of whole ChP 24 hours after saline or RBC injection.

(**F**) Brain section of E15.5 embryo 24 hours post-ICV injection of 70 kDa Texas Red dextran. Ventricle border = yellow dashed line; ChP outline = white dashed line. Epiplexus macrophages with dextran signal = gray arrows. Inset in red box. Scale bar = 100 µm. (**G**) Selected cells from ImageStream analysis positive for Texas Red dextran.

**Supplemental Figure S2**. (**A**) Proportion of ChP macrophages present in each cluster from Figure 4A. (**B**) Gene Ontology terms (Enrichr-KG)^71^ for the 3 non-replicating clusters of E16.5 macrophages. (**C**) FACS sorting of dual-labeled epiplexus (647, blue arrow) and stromal (Texas Red, red arrow) macrophages 3 hours following ICV/IP injection respectively from dissociated P7 ChP. (**D**) GPNMB and CSF1 staining in uninjected E14.5 mouse brain sections of the lateral ventricle and the ChP. Scale bar = 50 µm. (**E**) Violin plots of macrophage clusters and module score. (**F**) Dotplot of canonical macrophage markers compared to ChP macrophage clusters. (**G**) Comparison of bulk RNA sequencing genes from sorted stromal and epiplexus compared to BAM and YAM module scores. (**H**) Schematic and image of E13.5 choroid plaque. Scale bar = 200 µm. (**I**) Expression of common “marker” proteins in microglia/macrophages superior to the choroid plaque at E13.5. Scale bar = 200 µm. Box #1 and #2 show IBA1+ cells that co-express GPNMB, LYVE1, and IBA1 (white arrowheads). Scale bar = 100 µm.
