## Supplementary figures and images for "Fluid-Niche and Microglial Signatures Prime Robust Intraventricular Macrophage Response to Blood During Brain Development"

### Supplemental Figure 1

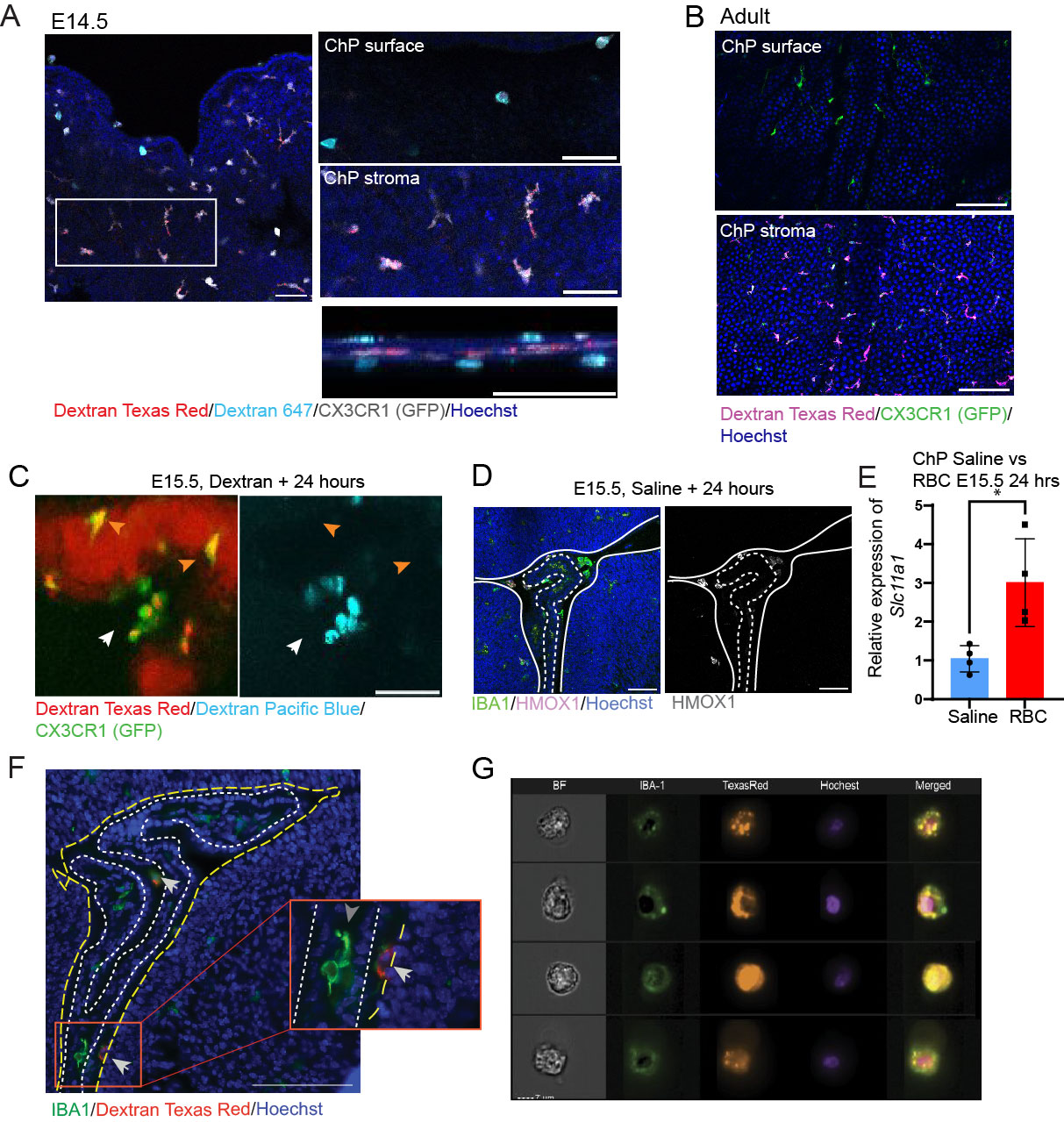

### Supplemental Figure 2

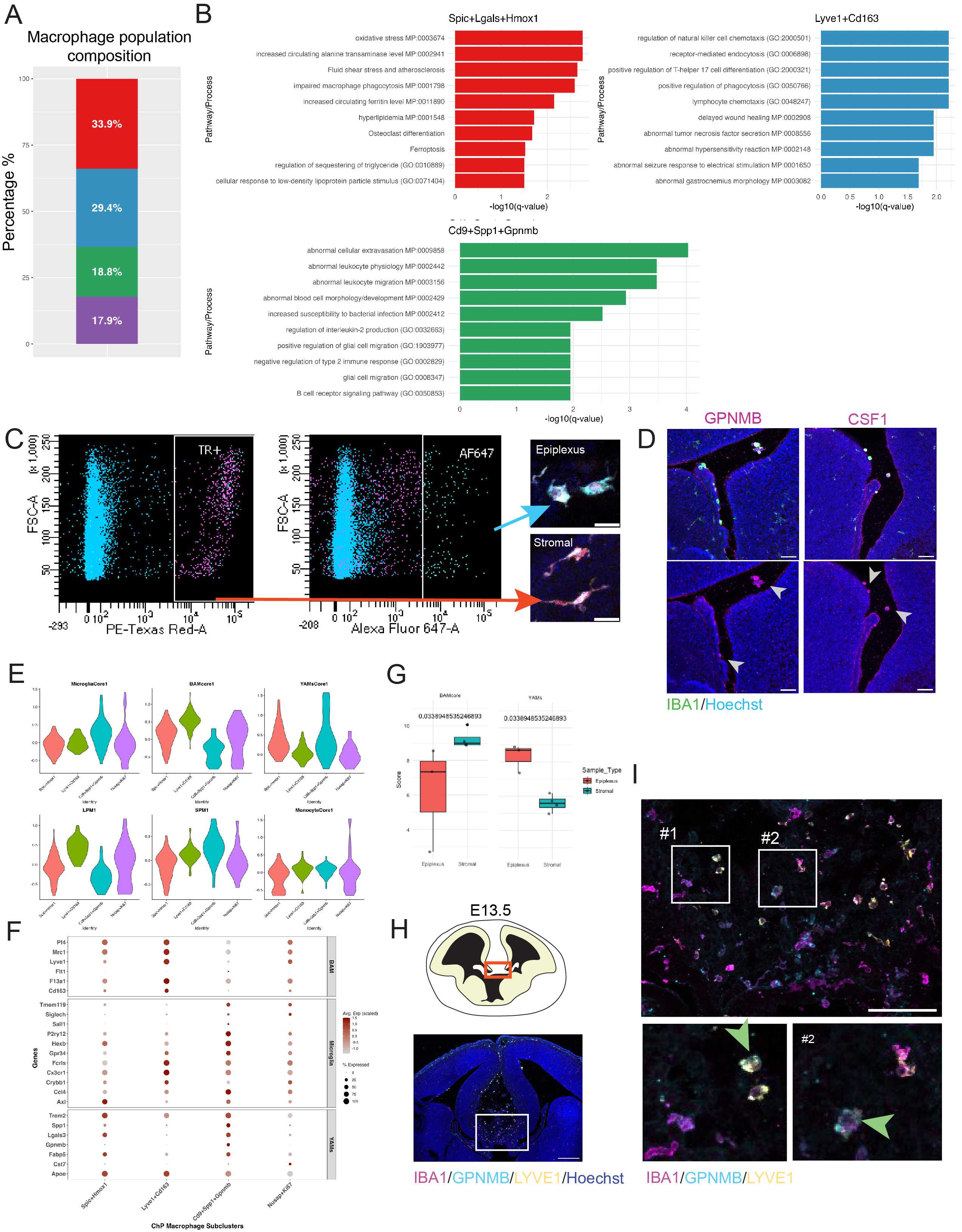
